# A combined EM and proteomic analysis places HIV-1 Vpu at the crossroads of retromer and ESCRT complexes: PTPN23 is a Vpu-cofactor

**DOI:** 10.1101/2021.02.22.432252

**Authors:** Charlotte A. Stoneham, Simon Langer, Paul D. De Jesus, Jacob M. Wozniak, John Lapek, Thomas Deerinck, Andrea Thor, Lars Pache, Sumit K. Chanda, David J. Gonzalez, Mark Ellisman, John Guatelli

**Affiliations:** Department of Medicine, University of California, San Diego School of Medicine and Veterans Affairs San Diego Healthcare System, La Jolla, California, USA; Infectious and Inflammatory Disease Center, Sanford Burnham Prebys Medical Discovery Institute, 10901 North Torrey Pines Road, La Jolla, California, USA; Department of Pharmacology, Skaggs School of Pharmacy and Pharmaceutical Sciences, University of California, San Diego, La Jolla, California, USA; National Center for Microscopy and Imaging Research, Center for Research on Biological Systems, University of California, San Diego, School of Medicine, La Jolla, California, USA; Department of Neurosciences, University of California, San Diego School of Medicine, La Jolla, California, USA

## Abstract

The HIV-1 accessory protein Vpu modulates membrane protein trafficking and degradation to provide evasion of immune surveillance. Targets of Vpu include CD4, HLAs, and BST-2. Several cellular pathways co-opted by Vpu have been identified, but the picture of Vpu’s itinerary and activities within membrane systems remains incomplete. Here, we used fusion proteins of Vpu and the enzyme ascorbate peroxidase (APEX2) to compare the ultrastructural locations and the proximal proteomes of wild type Vpu and Vpu-mutants. The proximity-omes of the proteins correlated with their ultrastructural locations and placed wild type Vpu near both retromer and ESCRT-0 complexes. Hierarchical clustering of protein abundances across the mutants was essential to interpreting the data and identified Vpu degradation-targets including CD4, HLA-C, and SEC12 as well as Vpu-cofactors including HGS, STAM, clathrin, and PTPN23, an ALIX-like protein. The Vpu-directed degradation of BST-2 required PTPN23 but not the retromer subunits. These data suggest that Vpu directs targets from sorting endosomes to degradation at multi-vesicular bodies via ESCRT-0 and PTPN23.

**Author Summary:** Vpu triggers the degradation or mis-localization of proteins important to the host’s immune response. Vpu acts as an adaptor, linking cellular protein targets to the ubiquitination and membrane trafficking machinery. Vpu has been localized to various cellular membrane systems. By fusing wild type Vpu and Vpu-mutants to the enzyme ascorbate peroxidase, we defined the cellular proteome in proximity to Vpu and correlated this with the protein’s location. We found that wild type Vpu is proximal to ESCRT proteins, retromer complexes, and sorting and late endosomal proteins. Functionally, we found that the Vpu-mediated degradation of the innate defense protein BST-2 required PTPN23, an ALIX-like protein, consistent with our observation of Vpu’s presence at the limiting membranes of multi-vesicular bodies.

## Introduction

HIV-1 encodes the accessory proteins Vif, Vpr, Nef, and Vpu to overcome cell-intrinsic, innate, and adaptive host defenses. Vpu is a small, non-enzymatic, integral membrane protein that functions as an adaptor, linking targeted cellular proteins to the protein quality control and membrane trafficking machinery to induce their re-localization or degradation. Cellular targets of Vpu interfere directly with viral replication or support immune surveillance; these targets include CD4 (the virus’s primary receptor), BST-2 (an interferon-induced protein that traps newly assembled virions on the infected-cell surface), natural killer (NK) cell receptors (NTB-A), class I MHC (HLA-C), CCR7, and tetraspanins (1–7).

Several Vpu cofactors and co-opted pathways have been identified. These include a Skp1/cullin1/F-box (SCF) multi-subunit E3 ubiquitin ligase containing β-TrCP (8). A phospho-serine acidic cluster (PSAC) in the cytoplasmic domain (CD) of Vpu binds β-TrCP, recruiting the E3 ligase and inducing poly-ubiquitination of certain Vpu-interacting proteins such as CD4 and BST-2. For CD4, ubiquitination precedes extraction from the endoplasmic reticulum (ER) and ultimate degradation by the proteasome (9). Other Vpu targets, such as BST-2, are instead degraded within the endo-lysosomal system (10, 11).

Vpu-mediated antagonism of BST-2 involves a complex interplay of membrane transport steps. Vpu retains newly synthesized BST-2 in the *trans*-Golgi network (TGN), and it inhibits the recycling of endocytosed BST-2 to the plasma membrane (12, 13). The net down-regulation of BST-2 from the cell surface requires clathrin and partly depends on the hetero-tetrameric clathrin adaptor (AP) complexes 1 and 2 (10, 14–16). An acidic leucine-based motif and the PSAC in Vpu’s CD bind the AP complexes directly, enabling Vpu to redirect the endosomal transport of its targets independently of the SCF E3 ubiquitin ligase (10, 15–18). Nonetheless, the endo-lysosomal degradation of BST-2 depends on Vpu’s PSAC as well as the SCF E3 ligase, and on the ubiquitin-binding protein HRS (HGS), a subunit of the ESCRT-0 complex (19).

The multifaceted mechanism of Vpu-action is reflected in the protein’s complex subcellular itinerary, which includes the ER, plasma membrane (PM), and various endosomal compartments. Consistent with this, microscopic data place Vpu at steady-state in the ER, Golgi and the TGN, the PM, and recycling endosomes (9, 20–22).

We asked whether a more integrated view of Vpu-activity could be obtained via a high-depth characterization of the protein’s proximal proteome. Proximity-labeling covalently tags protein-neighbors in living cells with small molecules such as biotin, enabling isolation by affinity-purification and identification and quantitation by mass spectrometry. This approach enables the identification of physiologically relevant interactions even when they are weak or transient. In the present study, we expressed Vpu fused to the enzyme APEX2, an ascorbate peroxidase whose catalytic activities enable both electron microscopic localization as well as the labeling of proximal proteins with biotin (23). The biotinylated proteins were captured, identified, and quantified using tandem-mass-tag (TMT)-based proteomics. To correlate ultrastructural localization with proximal proteins, we compared wild type Vpu to three Vpu-mutants: Vpu-A18H, a mutant of the Vpu transmembrane domain (TMD) that is retained in the ER; Vpu-AAA/F, a mutant of the alanine face of the Vpu TMD that interacts with the TMD of BST-2 and other Vpu targets; and Vpu-S52,56N, a mutant of the key serines of the PSAC motif, which is unable to interact with either β-TrCP or the medium (µ) subunits of AP-1 and AP-2 and is partly displaced from juxtanuclear endosomes to the plasma membrane.

Through these comparisons, we characterized the proximal proteome and itinerary of Vpu, including changes in response to the above substitutions in Vpu whose consequences for protein-interaction and function are partly known. Our results place wild type Vpu at sorting endosomes, from which its targets can be diverted via ESCRT-related proteins toward the interior of multivesicular bodies for degradation.

The results generate hypotheses regarding potential Vpu targets, such as SEC12, and they identify PTPN23 as a novel cofactor of Vpu.

## Results

### Vpu-APEX2 fusion is well expressed and functional

The enzyme ascorbate peroxidase 2 (APEX2) was genetically fused to the C-terminus of FLAG-tagged human codon-optimized clade B (NL4.3) Vpu (VpHu) with a short intervening linker sequence (Fig. 1A). APEX2 is modified for enhanced activity and improved detection sensitivity compared to its predecessor, APEX (23). Given the size of the tag (28 kDa) compared to that of Vpu (17 kDa), we first tested whether the fusion to APEX2 impaired Vpu activity. When transiently expressed in HeLa P4.R5 cells, which express the HIV receptors CD4, CXCR4, and CCR5 as well as BST-2, the Vpu(FLAG)-APEX2 fusions were well-expressed as measured by western blotting (Fig. 1B). Vpu-induced downregulation of surface BST-2 and CD4 was measured by immunofluorescent staining and flow cytometry (Fig.1). APEX2-tagged Vpu retained biological activity against BST-2 and CD4, although it was slightly less active than Vpu tagged only with a C-terminal FLAG-epitope (Fig. 1C).

**Figure 1.**
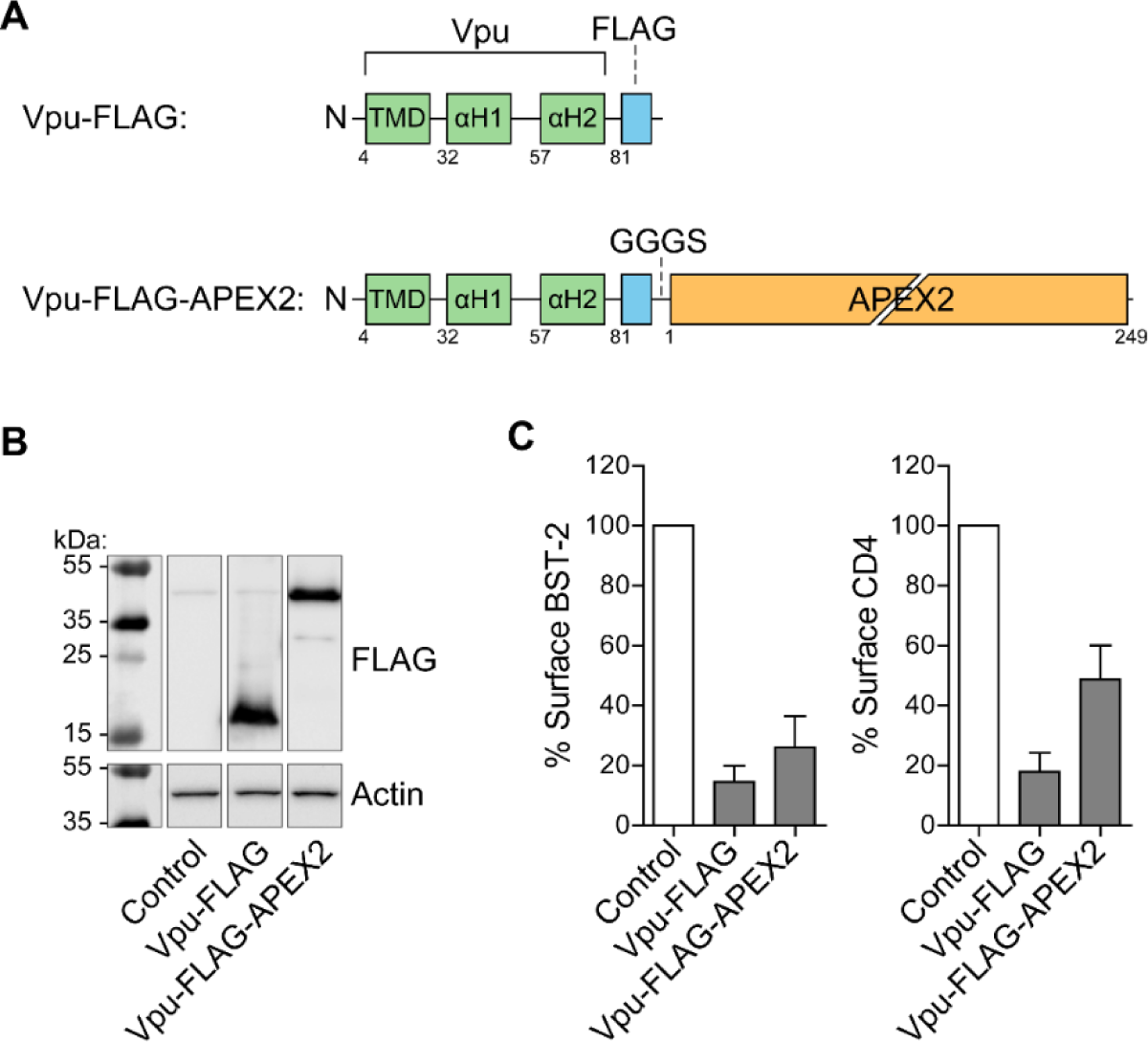
Vpu-APEX2 fusion protein design and activity. (A) A schematic representation of C-terminally tagged Vpu constructs; Vpu-FLAG and Vpu-FLAG-APEX2. A GGGS linker lies between the FLAG epitope and APEX2. (B) HeLa P4.R5 cells were transfected with Vpu constructs bearing C-terminal FLAG or FLAG and APEX2. Protein expression was analysed by western blot. (C) Cell-surface levels of BST-2 and CD4 were measured in the presence of Vpu-FLAG or Vpu-FLAG-APEX2 by flow cytometry. Surface levels of BST-2 and CD4 on cells expressing FLAG- or FLAG-APEX2-tagged Vpu was expressed as the % of control cells not expressing Vpu. Error bars represent standard deviation of n=3 experiments.

### Vpu-APEX2 distorts juxtanuclear endosomes and labels the limiting membranes of multi-vesicular bodies (MVBs)

APEX enables the generation of electron-dense material in fixed cells that can be detected by transmission electron microscopy (24). An advantage of APEX over horseradish peroxidase (HRP) is that it maintains activity in the reducing cytosolic environment. This allows APEX to be used for both intracellular protein imaging by electron microscopy and spatially resolved protein mapping (Martell, Deerinck et al. 2012).

The subcellular distribution of wild-type Vpu-APEX2 was first evaluated by electron microscopy (Fig. 2). HeLa cells were transiently-transfected to express Vpu-FLAG or Vpu-FLAG-APEX2. The next day, the cells were fixed and reacted with DAB in the presence of hydrogen peroxide (H_2_O_2_). After staining with osmium tetroxide, the polymerized DAB was visualized as a dark, electron-dense stain. This stain, corresponding to the location of Vpu-APEX, was observed on the cytoplasmic surface of vesiculated juxtanuclear membranes that appeared to be derived from the Golgi and endosomal systems (Fig. 2B). Most of these vesicles were unilamellar, enlarged vesicles (EVs), but some were consistent with multi-vesicular bodies (MVBs). The Vpu-APEX2 stain was restricted to the limiting membrane of MVBs and did not appear on intra-lumenal vesicles. Vesiculated Golgi was observed in cells transfected with Vpu-FLAG (without the APEX2 tag; Fig. 2C). In contrast, in non-transfected HeLa P4.R5 cells, the Golgi appeared as a typical stack of flattened cisternae (Fig. 2D). These data are consistent with Vpu’s known activities and residence in biosynthetic endo-lysosomal membranes (14, 15, 25, 26).

**Figure 2.**
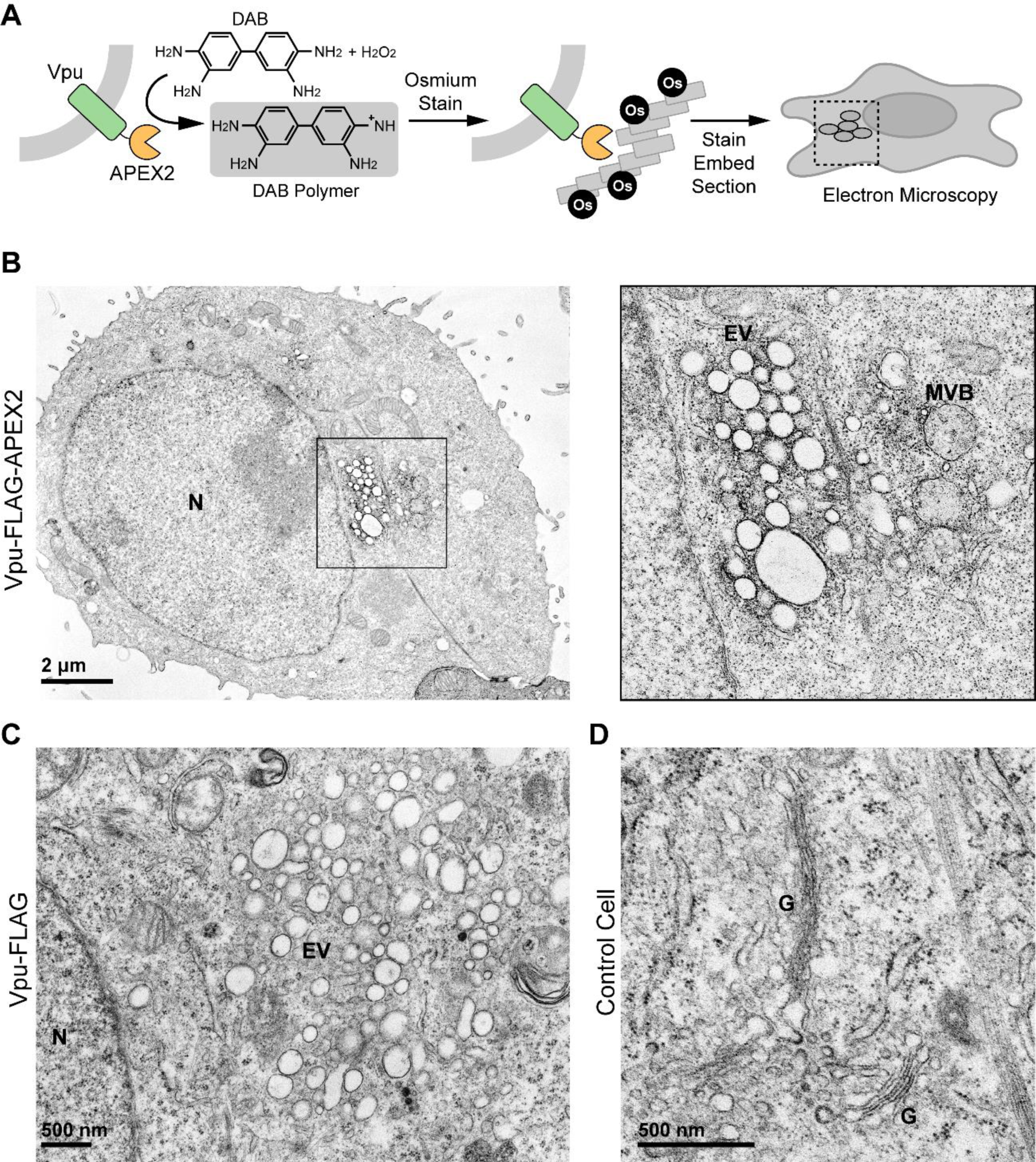
Vpu-APEX2 localizes to enlarged juxtanuclear endosomes and the limiting membranes of multi-vesicular bodies. (A) Schematic depicting APEX2 staining protocol for visualization by electron microscopy. (B) HeLa P4.R5 cells were transfected with the codon-optimized Vpu constructs bearing a C-terminal APEX2 tag. 24 hours later the cells were fixed before APEX2-dependent polymerization of DAB in the presence of hydrogen peroxide. The cells were then stained with OsO4, processed, and imaged by transmission electron microscopy (TEM). Left panel: Cells expressing Vpu-APEX2 contain juxtanuclear accumulations of osmium-highlighted enlarged vesicles (EV) likely derived from the Golgi and endosomes. Right panel: a higher magnification image of the juxtanuclear region of the cell shown at left. The limiting membranes of vesicles resembling multivesicular bodies (MVB) are highlighted by osmium. (C) Cells expressing Vpu (without an APEX2 tag) also contain enlarged juxtanuclear vesicles. (D) A control image showing Golgi (G) stacks in non-transfected HeLa P4.R5 cells.

### Localization of Vpu-APEX2 mutants: Vpu-A18H mis-localizes to the ER; Vpu-S52,56N mis-localizes to the plasma membrane

We then compared the light and electron microscopic localization of Vpu-FLAG with Vpu-FLAG-APEX2 when both constructs contained previously characterized mutations. We evaluated three Vpu-mutants: Vpu-A18H, a mutant of the Vpu transmembrane domain (TMD) that is retained in the ER (26); Vpu-AAA/F, a mutant of the alanine face of the Vpu TMD that interacts with the TMD of BST-2 and potentially other Vpu targets (27); and Vpu-S52,56N, a mutant of the key serines of the PSAC motif (8). HeLa P4.R5 cells transfected to express Vpu WT or the Vpu mutants tagged with FLAG were fixed and stained for the FLAG epitope and the *trans* Golgi resident marker protein TGN-46, then visualized by immunofluorescence microscopy (Fig. 3A). Vpu-FLAG localized to juxtanuclear membranes and overlapped partially with TGN-46, in agreement with previous studies (15, 20, 22). As anticipated, Vpu A18H was restricted to the nuclear envelope and a cytoplasmic, ER-like distribution. The AAA/F mutation did not alter the localization of Vpu appreciably. In contrast, Vpu-S52,56N displayed a relatively more dispersed cytoplasmic staining.

**Figure 3.**
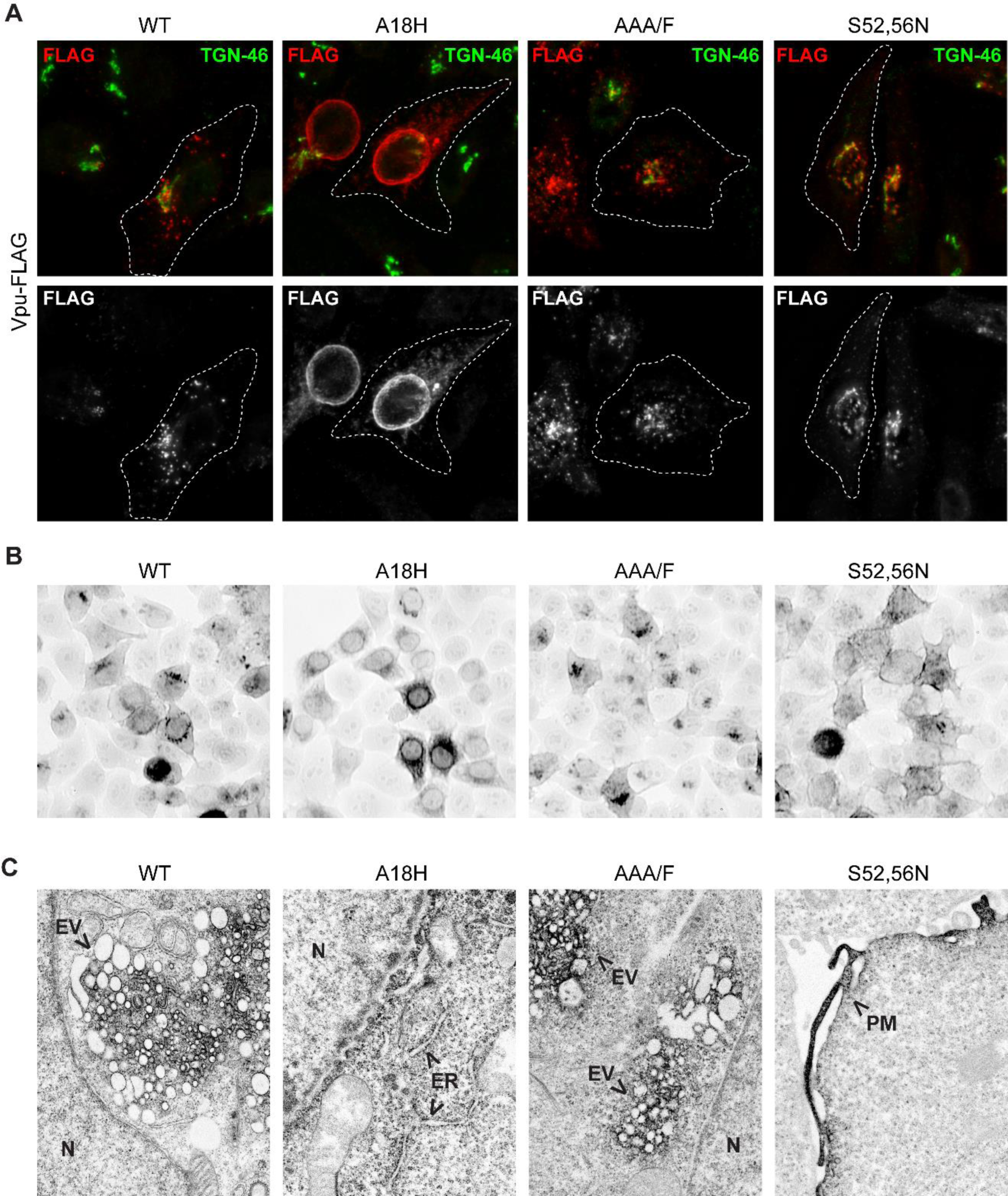
Light and electron microscopic distributions of Vpu-APEX2 and mutants A18H, AAA/F, and S52,56N. (A) HeLa P4.R5 cells were transfected to express Vpu-FLAG (A), either wild type or encoding the mutations A18H, AAA/F, or S52,56N. The cells were fixed and stained for FLAG (shown in red) and TGN-46 (shown in green). (B) HeLa P4.R5 cells were transfected to express Vpu WT and the indicated mutants tagged with APEX2; 24 hours later the cells were reacted with DAB in the presence of hydrogen peroxide. The cells were stained and osmiophilic DAB polymer visualized in whole-cells by brightfield microscopy. (C) Thin-section electron microscopy of cells expressing Vpu WT-, A18H-, AAA/F-, or S52,56N-APEX2. Arrows indicate concentrations of osmiophilic polymer stain. N = nucleus, ER = endoplasmic reticulum, PM = plasma membrane.

Similar distributions were observed at the light microscopic level when Vpu-APEX2 and the mutants were visualized using H_2_O_2_, DAB, and osmium (Fig. 3B). The distribution of WT Vpu-APEX2 was juxta- and peri-nuclear, whereas Vpu-A18H-APEX2 was ER-like. In contrast, Vpu-S52,56-APEX2 outlined the cell perimeter, consistent with residence at the plasma membrane. Electron microscopic images (Fig. 3C) revealed vesiculation of juxtanuclear membranes by the AAA/F and S52,56N mutants as well as by the wild type Vpu-APEX2 (Fig. S1). Vpu-A18H-APEX2 stained the nuclear envelope and ER membranes and at high levels of expression induced a striking alteration in the structure of these organelles (Fig. S2). Vpu-S52,56N-APEX2 stained the plasma membrane, consistent with the light-microscopic observations. These data indicated that the subcellular distribution of Vpu-APEX2 was similar to that of Vpu without the APEX-tag. The data also indicated that specific mutations in Vpu-APEX2 modulated the protein’s subcellular distribution in a manner consistent with known properties of Vpu.

### The Vpu-proximity-ome defined by multiplexed quantitative proteomics and comparison of Vpu mutants

In living cells and in the presence of hydrogen-peroxide, APEX can catalyze the generation of biotin-phenoxy radicals to enable rapid, spatially restricted labeling of proximal proteins; these proteins can then be isolated by standard pull-down methods and identified by mass spectrometry. An advantage of APEX-based proximity labeling is that weak interactions can be identified, which would otherwise be lost during standard affinity purification. Moreover, neighboring proteins should be labeled and identified, even if they are not direct interactors. To evaluate the feasibility of this approach in the case of Vpu, we first detected proteins biotinylated by Vpu(FLAG)-APEX2 using immunofluorescence and immunoblot.

HeLa P4.R5 cells were transiently-transfected to express Vpu(FLAG)-APEX2 or the previously-characterized Mito-Matrix-APEX2 (23). The next day, the cells were incubated with hydrogen peroxide in the presence of biotin-phenol for 1 minute, fixed, and stained with streptavidin conjugated to the fluorescent label Alexa Fluor 594 (Fig. 4B). In cells transfected to express Vpu-APEX2, streptavidin was concentrated in perinuclear regions and overlapped with FLAG, consistent with biotinylation of proteins in close proximity to Vpu-APEX2, likely including Vpu-APEX2 itself. The streptavidin signal was also present faintly and diffusely throughout the nucleus and cytoplasm. In contrast, in cells transfected to express Mito-APEX2, streptavidin was restricted to mitochondrial structures without a diffuse background. This restriction presumably reflects the generation of biotin-phenoxy radicals within the enclosed membranes of mitochondria in the case of Mito-APEX2 rather than in the cytosol in the case of Vpu-APEX2. In a second set of experiments, cells expressing Vpu(FLAG)-APEX2 and related mutants, or expressing Mito-APEX2, were incubated with hydrogen peroxide in the presence of biotin-phenol for 1 minute, lysed, and the proteins were separated by SDS-PAGE and analyzed by western blot. Biotinylated proteins were detected using streptavidin conjugated to HRP. The size distribution of proteins biotinylated by Vpu-APEX2 was strikingly different from that of Mito-APEX2, but the distribution of proteins biotinylated by WT Vpu-APEX2 and the related Vpu-mutants were indistinguishable. For the Vpu proteins, the most abundant band at 40 kDa likely corresponds to self-biotinylation of Vpu-APEX2. Minimal background biotin labeling was observed in cells transfected with empty plasmid (control) or Vpu(FLAG) lacking the APEX2 tag.

**Figure 4.**
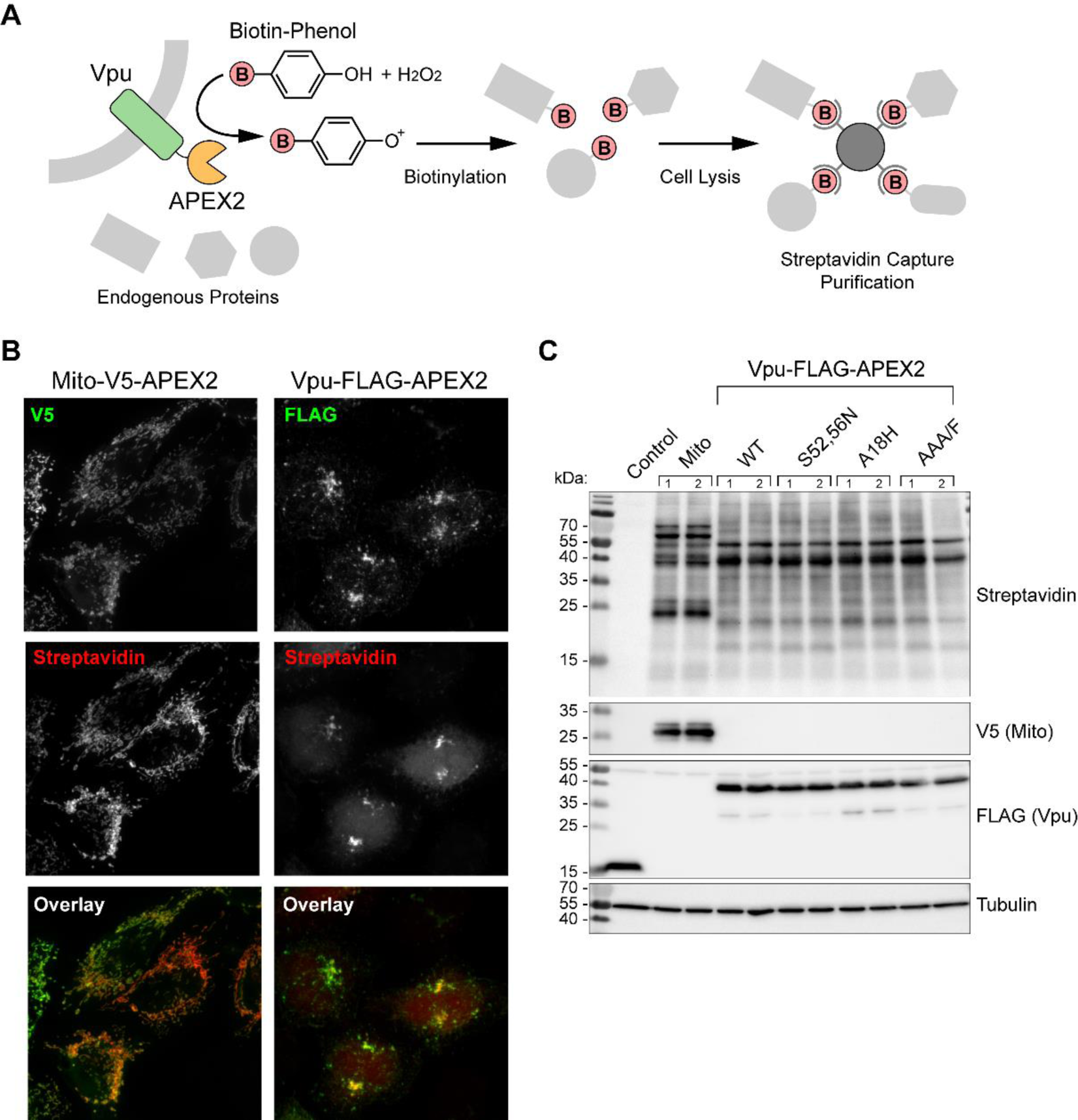
Biotinylation of Vpu-APEX2 proximal proteins. (A) Schematic representation of APEX2-mediated biotinylation reaction; in the presence of hydrogen peroxide and biotin-phenol, APEX2 catalyses biotinylation of nearby proteins, which can be captured by streptavidin. (B) Detection of biotinylated proteins by immunofluorescent-streptavidin. HeLa P4.R5 cells expressing Mito-V5-APEX2 or Vpu-FLAG-APEX2 were incubated with biotin phenol and hydrogen peroxide for 1 minute before quenching, fixation, and staining with streptavidin-alexa594 (red) and anti-V5 or anti-FLAG (green). (C) Biotinylation pattern of protein species visualized by western blot; HeLa P4.R5 cells transfected to express Mito-V5-APEX or WT Vpu-, S52,56N-, A18H-, or AAA/F-FLAG-APEX2 were incubated with biotin-phenol, lysed and proteins separated by SDS-PAGE and western blot. Biotinylated proteins were detected using streptavidin-HRP. The control cells were transfected to express Vpu-FLAG only; streptavidin staining is absent in the absence of APEX2.

For our preliminary mass spectrometry experiments, we transfected HeLa P4.R5 cells to express the WT Vpu-APEX2, or the mutants, A18H, AAA/F, S52,56N, or the Mito-APEX2 control. The next day, the APEX2-catalyzed biotinylation procedures were performed, and biotinylated proteins were captured on streptavidin-coated beads. The captured proteins were eluted from the beads and analyzed by quantitative proteomics. Data were normalized as described in the Materials and Methods and are presented as a heatmap of the relative abundance of 1779 common proteins identified in all samples (Fig. S3A). As anticipated, the Mito-APEX2 induced biotinylation of mitochondrial proteins, reflected in the high relative enrichment of proteins which conform to the Gene Ontology (GO) term “mitochondrial matrix” and other mitochondrion-associated cellular components (Fig. S3B, S3C).

In two subsequent independent experiments, the Vpu WT and mutants were compared for differential enrichment of biotinylated proteins that might represent potential cofactors or targets (Fig. 5 and Fig. 6). The data were first compared by pair-wise analysis of Vpu WT vs. individual mutants (Fig. 5). Relaxed statistical parameters were used to compare the relative enrichment of proteins between Vpu WT and the mutants, as the fold changes were relatively low; data are presented as the relative protein enrichment in the presence of Vpu mutants and wild-type Vpu, with *p*-value determined by *t*-test. The proximity-ome of the A18H mutant was enriched for proteins derived from the biosynthetic pathway (such as SEC proteins) and COPII-coated vesicles, while depleted in plasma membrane and endosomal proteins relative to that of wild type Vpu. These data were consistent with the ultrastructural imaging, which demonstrated restriction of the A18H mutant to predominantly the ER and nuclear envelope. The AAA/F mutant, whose ultrastructural distribution was similar to that of the WT Vpu, had few proteins significantly enriched or depleted relative to the WT protein (data not shown). In contrast, the proximity-ome of the S52,56N mutant, which at the light and EM level was partially redistributed to the plasma membrane, was significantly enriched in plasma membrane proteins including EGFR and known targets of Vpu, (CD4, CD81, and HLA-C), while depleted in endosomal proteins relative to that of wild type Vpu.

**Figure 5.**
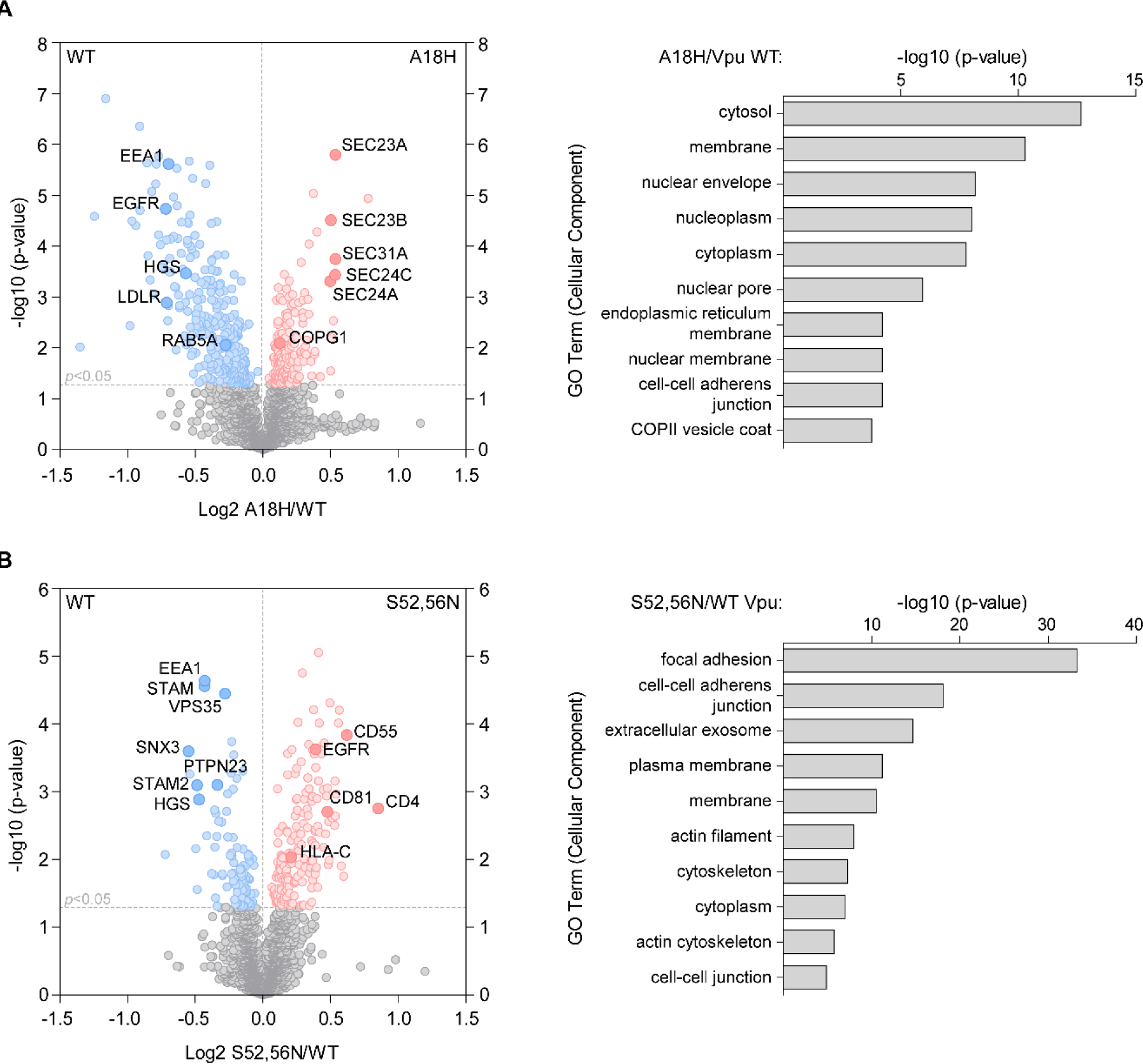
Pair-wise comparisons of the proximity-omes of wild type Vpu relative to the ER-retained Vpu-A18H and the plasma-membrane-enriched Vpu-S52,56N mutants. HeLa P4.R5 cells were transfected to express Vpu constructs bearing C-terminal APEX2 tags, in duplicate. Following proximity biotinylation reactions, the biotinylated proteins were isolated and subject to quantitative mass spectrometry. Volcano plots of protein enrichment in the presence of Vpu mutants A18H or S52,56N relative to the wild-type Vpu are shown, n=2 experiments. Significantly enriched proteins highlighted in red and blue are derived from the Student’s *t*-test (*p*<0.05). (A) The A18H mutant was enriched in proteins derived from the biosynthetic pathway, including the ER (labeled), while WT Vpu was enriched for proteins associated with plasma and endosomal membranes (labeled). (B) The S52,56N mutant was enriched in proteins derived from the plasma membrane (labeled), while WT Vpu was again enriched in endosomal sorting proteins (labeled). Proteins enriched by the S52,56N mutation included the known targets CD4, CD81, and HLA-C, and possible targets EGFR and CD55. For both (A) and (B), the x-axis shows log2 fold change of proteins enriched by mutant/WT Vpu and the y-axis -log10 of *p*-value derived from Student’s *t*-test. The 10 most highly enriched gene ontology (cellular component) terms are shown on the right of each volcano plot, corresponding to significantly enriched proteins proximal to the mutants; *p*-value derived from Bonferroni test.

**Figure 6.**
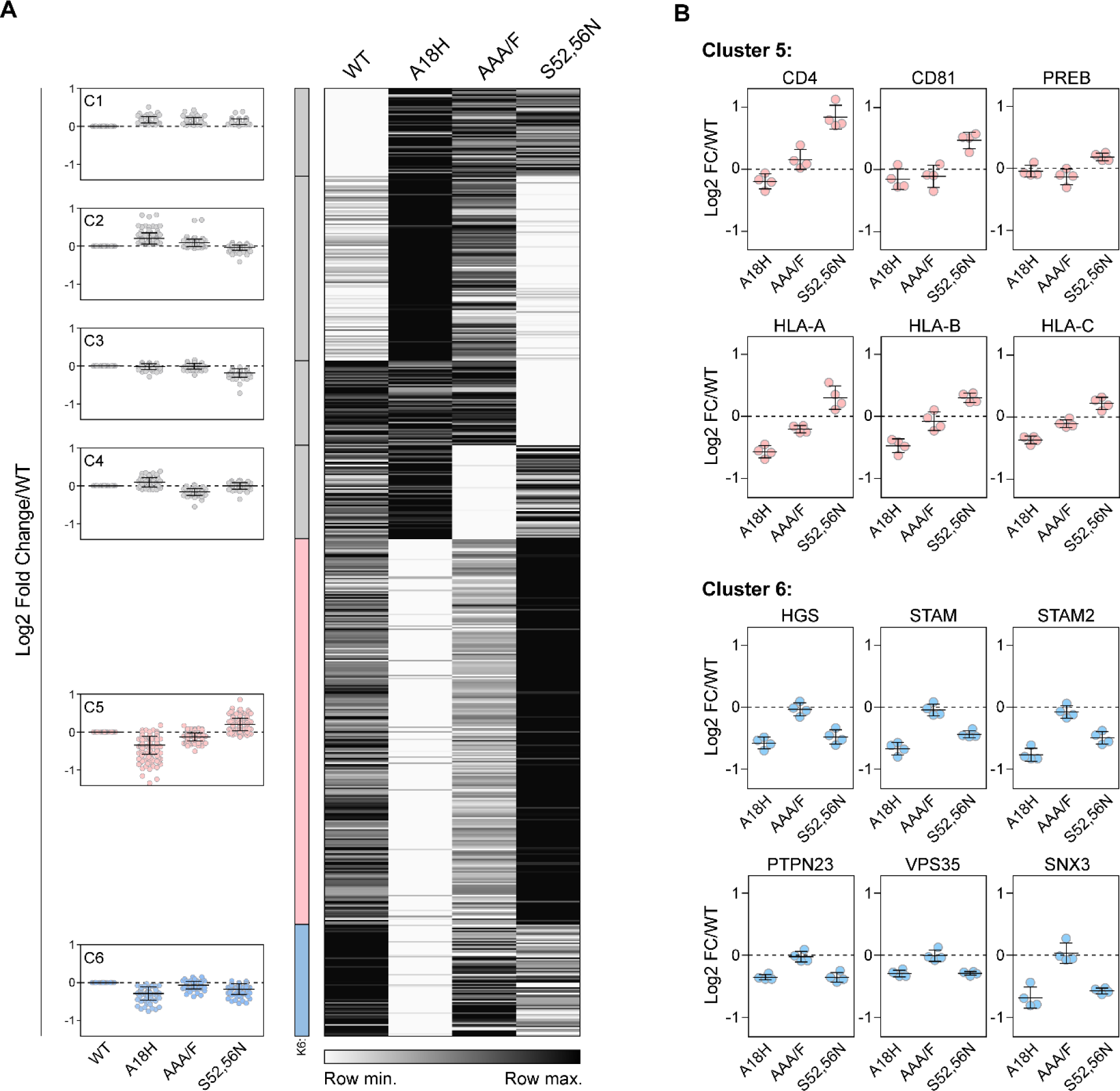
Heat map and k-means clustering of proteins (384) for which any Vpu-mutant was significantly different from wild type. (A) Heatmap of the relative abundance of proteins measured in Vpu-APEX2 WT and mutant samples. The heatmap was sorted into 6 clusters by k-means clustering analysis; the cluster profile is shown on the left. Data are presented as the fold change in protein abundance relative to WT. (B) Cluster 5 contains known and potential targets of Vpu, including CD4, CD81, and HLA-C. k-means cluster 6 contains potential serine-dependent cofactors of Vpu, including endosomal sorting proteins. Data are derived from duplicate samples per condition, from two independent experiments.

K-means clustering analysis of the patterns of protein enrichment across the four conditions (WT Vpu and the three mutants) highlighted potential cofactors (cluster 6) and targets (cluster 5) for Vpu activities (Fig. 6.) Specifically, we reasoned that cluster 5 proteins, which were decreased relative to WT when Vpu was retained in the ER by the A18H mutation but increased when Vpu was displaced to the plasma membrane by the S52,56N mutation, would include Vpu targets, especially if degraded by an ERAD-like mechanism. Consistent with this notion, cluster 5 includes known Vpu targets CD4, CD81, ICAM-1, and HLA proteins. We also reasoned that cluster 6 proteins, which were decreased by both the A18H mutation and the S52,56N mutation, could reflect the local intracellular environment of WT Vpu. Cluster 6 includes a variety of endosomal sorting proteins including the early endosomal protein EEA1, components of the ESCRT-0 complex (HGS, STAM, and STAM2), and components of the retromer complex (Vps35 and SNX3), among others. These data place wild type Vpu at the sorting endosome, a crossroads in the endosomal system at which Vpu is well-positioned to inhibit recycling and target proteins to endo-lysosomal degradation.

### Potential novel Vpu targets

We reasoned that proteins enriched in proximity to Vpu-S52,56N relative to WT Vpu could also represent proteins that are stabilized when Vpu is mutated and might be targets of Vpu-mediated degradation. To distinguish such targets of Vpu-mediated degradation from proteins that instead reflect changes in the local environment of Vpu induced by the displacement of the S52,56N mutant to the plasma membrane, we first measured surface-downregulation using flow cytometry. HeLa P4.R5 cells were transfected to express Vpu-FLAG, and surface EGFR, CD55, and CD4 were measured (data not shown). CD4 was downregulated by Vpu as expected, but EGFR and CD55 were unaffected; these proteins likely become more proximal to Vpu when Vpu is displaced to the plasma membrane and are unlikely to be a degradation-targets.

To screen further for Vpu targets, we used the relatively high throughput Global Arrayed Protein Stability Analysis (GAPSA) assay (28). cDNAs encoding approximately 160 proteins that were significantly enriched in proximity to the S52,56N mutant relative to WT Vpu were screened for degradation by WT Vpu, using Vpu-S52,56N as a control. We found eight proteins that were significantly degraded by WT Vpu but not by the S52,56N mutant (Fig. 7). Degradation of these proteins was confirmed by western blotting of the V5-tagged target proteins. Three of these proteins, CD4, HLA-C, and CD99, are known Vpu targets, but the others, including PREB/SEC12, are newly identified.

**Figure 7.**
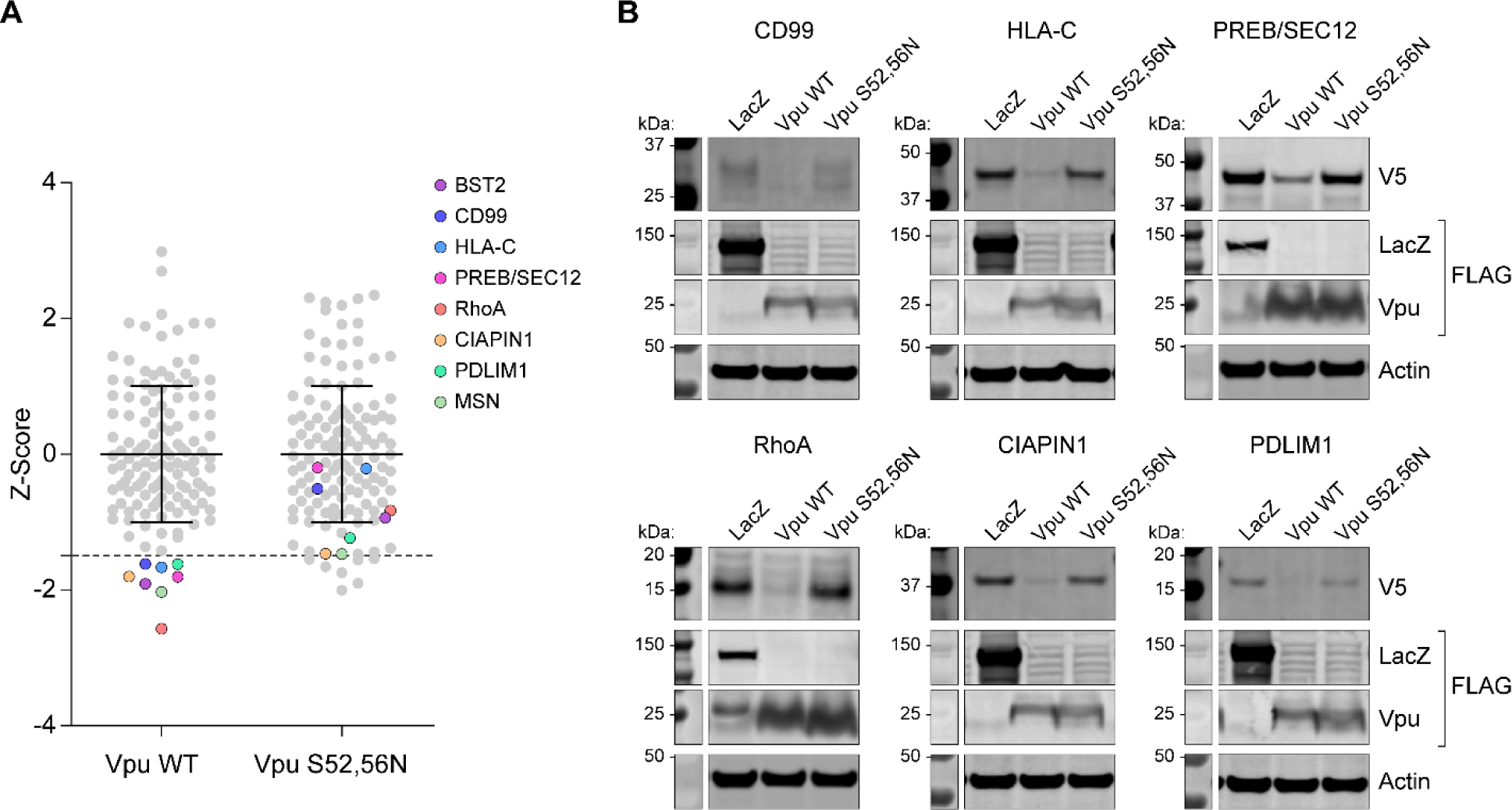
Proteins enriched in the proximity-ome of Vpu-S52,56N relative to wild type: Vpu-mediated and serine-dependent decreases in steady-state expression. (A) HEK293T cells were co-transfected with target cDNAs and plasmids expressing WT Vpu or Vpu-S52,56N. Levels of proteins were measured 48 hours later by immunofluorescent staining and automated quantification of fluorescence. Candidate Vpu targets whose expression is significantly decreased by Vpu are colored. (B) Vpu-mediated degradation of putative target proteins was tested by western blotting of exogenously-expressed cDNA targets.

### Potential novel Vpu cofactors

We reasoned that proteins enriched in proximity to WT Vpu relative to Vpu-S52,56N might represent cellular cofactors, as the DS52GxxS56 sequence has been shown to be a relatively promiscuous motif with regard to recruiting both the SCF-E3 ligase and clathrin AP complexes (8, 16, 18). STRING analysis of proteins identified in k-means cluster 6, which includes proteins whose proximity to Vpu is decreased by the S52,56N substitution, revealed a network of interrelated proteins involved in ubiquitin sorting, endosomal trafficking, and retromer-mediated trafficking (Fig. 8A).

**Figure 8.**
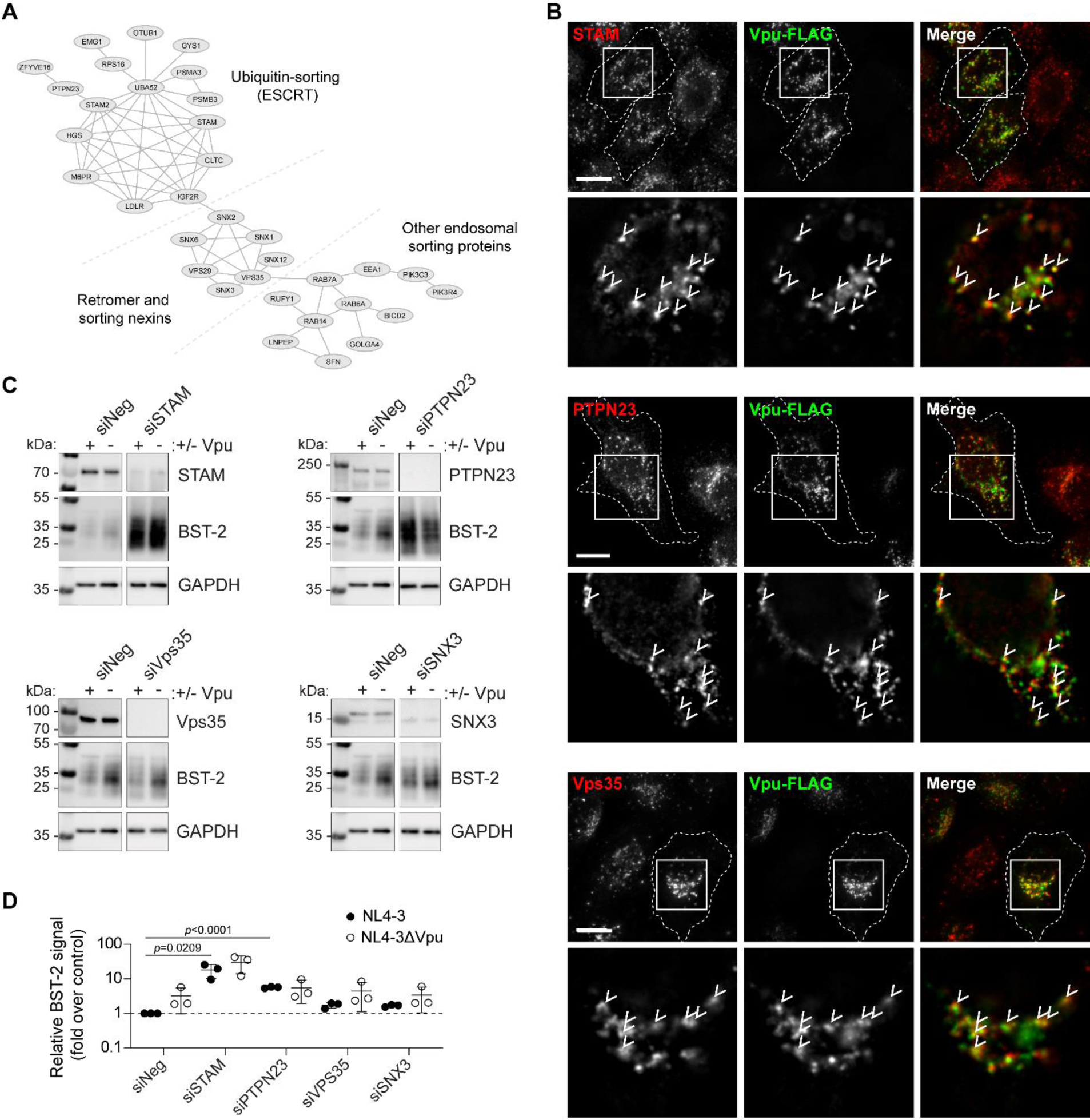
Potential Vpu cofactors: STRING relationships, co-localization with Vpu, and effects on expression of the Vpu-target BST-2. (A) The interactions of proteins in k-means cluster 6 (Figure 6) were visualized using the STRINGdb (Cytoscape) network analysis tool. The interrelated proteins identified represent candidate Vpu cofactors. (B) Immunofluorescence microscopy of Vpu-FLAG and some of the candidate cofactors. HeLa P4.R5 cells were transfected to express Vpu-FLAG. Cells were fixed and stained for the indicated endogenous proteins. Images are z-stack projections of full cell volumes; insets show single z-sections, with arrows indicating colocalized foci. Scale bars are 10 µm. (C) Candidate cofactor proteins were transiently knocked-down using siRNAs in HeLa P4.R5 cells. The cells were transfected with pNL4-3 (an HIV proviral plasmid expressing the complete viral genome including Vpu) or pNL4-3ΔVpu 48 hours later. The cells were lysed 24 hours post-transfection and BST-2 was probed by SDS-PAGE and western blotting. (D) BST-2 signals were measured relative to loading control (GAPDH) and presented as fold signal over NL4-3 (negative control siRNA; Vpu-expressed). Data are mean +/-SD of three independent experiments; *p*-value determined by Student’s *t*-test.

We investigated whether Vpu-FLAG colocalized with these proteins by immunofluorescence microscopy. HeLa P4.R5 cells were transfected to express Vpu-FLAG then fixed and stained for FLAG and either STAM, PTPN23, or Vps35; each colocalized with Vpu (Fig. 8B). We also observed partial colocalization between Vpu and the retromer protein SNX3 (Fig. S4).

To examine whether these proteins have roles in Vpu activities, we tested the ability of Vpu to degrade BST-2 in their absence. HeLa P4.R5 cells were depleted of STAM, Vps35, SNX3, or PTPN23 using siRNAs before transfection with proviral plasmids encoding either wild type HIV-1 clone NL4-3 (encoding Vpu) or NL4-3ΔVpu. Total cellular BST-2 was measured by western blotting 24 hours after transfection of the proviral plasmids (Fig. 8C). As expected, NL4-3 encoding Vpu reduced the levels of BST-2 compared to NL4-3ΔVpu in cells treated with the control siRNA (siNeg) (Fig. 8C). Knockdown of the ESCRT-0 subunit STAM markedly increased the steady-state levels of BST-2, but Vpu still reduced those levels. Knockdown of PTPN23, an ALIX-like protein involved in the formation of intralumenal vesicles in MVBs (29), inhibited the Vpu-mediated degradation of BST-2 without substantially affecting BST-2 levels in the absence of Vpu. Depletion of the retromer proteins Vps35 and SNX3 had minimal if any effect on Vpu-mediated degradation of BST-2. These data support distinct roles for ESCRT-0 and PTPN23 in the degradation of BST-2: whereas the ESCRT-0 protein STAM supported physiologic degradation of BST-2 in the absence of Vpu, PTP23N instead supported Vpu-mediated degradation specifically, consistent with activity as a Vpu cofactor.

## Discussion

Proteomic methods have been used to explore host-HIV protein-protein interactions and to look broadly at changes in the plasma membrane induced by the virus (30, 31). Here, we used proximity labeling mediated by APEX2 fusion proteins as a novel approach to define the proteome proximal to the HIV-1 protein Vpu, a membrane protein that mediates viral evasion of innate and adaptive immunity. The use of Vpu mutants whose subcellular localization differed was key to distinguishing proteins specific to the neighborhood of wild type Vpu from a large background. We observed that Vpu-APEX2 (like Vpu without the APEX2 fusion-tag) distorted juxtanuclear endosomes, replacing typical stacks of thin Golgi cisternae with enlarged vesicles. These vesicles, as well as the limiting membranes of MVBs, were labeled when cells expressing Vpu-APEX2 were visualized by thin section electron microscopy. When Vpu was mutationally trapped in the ER, its proximal proteome included an abundance of ER-associated proteins including components of COPII coats. When Vpu was mutationally displaced to the plasma membrane, its proximal proteome was depleted of early and sorting endosomal components; these included subunits of the retromer and the ESCRT-0 complexes, as well as the ALIX-like protein PTPN23, which supported the degradation of BST-2 by Vpu. Comparison of the proximal proteomes of the wild type and mutant Vpu proteins yielded a list of proteins up-regulated by substitution of the serines within the protein’s PSAC motif. Some of these are likely proximity markers of the plasma membrane, consistent with the Vpu mutant’s change in localization, while others, including HLA-C, CD99, and SEC12, are potentially subject to Vpu-directed degradation, a possibility supported by the transient expression experiments herein.

Many proteins identified here are unlikely to be either Vpu targets or cofactors, but nonetheless contribute to creating a picture of Vpu’s itinerary within cellular membranes. That itinerary seems focused on sorting endosomes identified by the specific presence of EEA1 and ESCRT-0 subunits in the neighborhood of wild type Vpu. However, it also likely includes late endosomes and MVBs, consistent with the presence of the ALIX-like protein PTPN23, which supports the budding of intralumenal vesicles into MVBs (29). This characterization of Vpu is reinforced by the labeling of the limiting membranes of MVBs by wild type Vpu-APEX2 when the enzyme is used to generate an osmiophilic reaction product.

PTPN23 is required for the sorting of EGFR into MVBs and its ultimate degradation (32). Consistent with this, functional data herein indicate that PTPN23 is a cofactor of Vpu on the path of endo-lysosomal degradation of at least one of its targets, BST-2. Whether PTPN23 directly interacts with Vpu remains an open question. Nonetheless, PTPN23 does not seem required for the physiologic degradation of BST-2, suggesting that it defines a distinct pathway of degradation co-opted by Vpu. Although not found in the heatmap of Figure 6 (see Supplemental Table 2), the PTPN23-associated protein CHMP 4B and the deubiquitinase USP8/UBPY co-clustered with PTPN23 and STAM in the independent experiments shown in supplemental figure S2 (see Supplemental Table 1)(32). These data are consistent with a model in which Vpu co-opts ESCRT-0, PTPN23, and CHMP 4B to direct targets into the lumen of MVBs for degradation.

The ESCRT-0 components HRS and STAM were identified as proximal to wild type Vpu, but in contrast to PTPN23, knockdown of STAM markedly increased the expression of BST-2 in either the absence or presence of Vpu. This suggests that ESCRT-0, which plays a key role in the sorting of ubiquitinated membrane proteins from early endosomes to late endosomes at the expense of recycling to the plasma membrane (33), constitutively targets BST-2 toward endo-lysosomal degradation. Vpu presumably acts downstream of that step, since it can stimulate partial degradation of BST-2 even when the expression of BST-2 is increased by knockdown of STAM.

Although our primary intention was not to look for new Vpu-targets, our data suggest several novel targets of serine-dependent Vpu-mediated degradation. One of the more intriguing is PREB, also known as SEC12, a guanine nucleotide exchange factor for the GTPase Sar1p (34), which regulates the formation of COPII coats and ER-to-Golgi transport. Degradation of SEC12 by Vpu, shown here in transient expression experiments, could cause a block in ER-to-Golgi transport and is potentially consistent with the reported inhibition of exocytic membrane trafficking by Vpu (12, 35). Although it might also be consistent with the formation of the large juxtanuclear endosomes observed electron microscopically, those structures were not serine-dependent (Fig. S1). On the other hand, cells expressing the ER-restricted Vpu-A18H mutant often showed exuberant accumulation of ER membranes emanating from the nuclear envelope, potentially consistent with an exaggerated SEC12 degradation phenotype and a block in ER-to-Golgi transport (Fig. S2).

In summary, correlative microscopic and proteomic analyses have provided a view of the Vpu-proximal proteins with unprecedented depth. The data place wild type Vpu predominantly at early sorting endosomes as well as at late endosomes and MVBs. The data generate new models, including the role of PTPN23 in degradation directed by Vpu at the MVB and the possibility that Vpu-mediated degradation of SEC12 underlies inhibition of exocytic trafficking. Elaborating these new models will require viral expression of Vpu in natural host cells such as CD4-positive T cells or macrophages.

## Materials and Methods

### Cells

HeLa P4.R5 cells, which express the HIV-1 receptors CD4 and CCR5, were obtained from the NIH AIDS Research and Reference Reagent program from Dr. Nathaniel Landau (36). HEK293 cells were obtained from Dr. Saswati Chaterjee (City of Hope). HEK293T cells (used in GAPSA assays) were purchased from ATCC (Manassas, VA). All cell lines were maintained in Dulbecco’s modified Eagle medium (DMEM) supplemented with 10% fetal bovine serum (FBS), penicillin/streptomycin, and 1 µg/mL puromycin in the case of HeLa P4.R5 cells.

### Plasmids

The C-terminally FLAG-tagged human codon-optimized clade B (NL4.3) Vpu (VpHu) has been previously described (17). The pCG-GFP reporter plasmid (37) was provided by Dr. Jacek Skowronski, Case Western Reserve University, Cleveland, OH. pcDNA3.1-Connexin43-GFP-APEX2 construct was obtained from Addgene, deposited by Dr. Alice Ting, Massachusetts Institute of Technology, Cambridge, MA (23). The pcDNA3.1-based Vpu-FLAG-APEX2 was generated by overlap extension PCR amplification before restriction digest and ligation into the pcDNA3.1(-) plasmid backbone between NheI and EcoRI sites. Vpu-FLAG-containing fragment was amplified using 5’ AGATTCGCTAGCATGGTGCCCATTATTGTCGC and 5’ CACAGTTGGGTAAGACTTTCCGGAGCCGCCGCCCTTATCGTCGTCATCCTTGTAA primers, and FLAG-APEX2 was amplified using 5’ TTACAAGGATGACGACGATAAGGGCGGCGGCTCCGGAAAGTCTTACCCAACTGTG and 5’TGCTTAGAATTCTTAGGCATCAGCAAACCCAAG. Vpu-FLAG plasmid constructs encoding the mutations AAA/F, A18H and S52,56N were previously generated. The overlap extension PCR method was used to amplify and ligate these mutated DNAs into the pcDNA3.1(-) backbone with a C-terminal APEX2 tag. The Mito matrix-v5-APEX2 construct was generated in the Ting lab (23). Expression plasmids containing V5-tagged cDNAs in the pLX304 backbone were obtained from the Lenti ORFeome Collection (38). A pcDNA4-Vpu-FLAG plasmid was used for the GAPSA assay; LacZ-FLAG was used as a control (28). pcDNA4-Vpu-S52,56N-FLAG was generated by G-block synthesis (Integrated DNA Technologies, IDT) and ligation between BamHI and NotI sites of the pcDNA4 backbone, using In-fusion cloning reagent (Takara Bio). Full-length HIV-1 and HIV-1 lacking Vpu were expressed from the HIV-1 proviral plasmid pNL4-3 (39) and pNL4-3ΔVpu (40).

### siRNAs

The siRNAs targeting STAM and Vps35 were custom synthesized by Sigma-Aldrich, the target sequences were as follows; STAM: UAACUUGGUAUAUAAGGAAAGGGCC, and Vps35: GCCUUCAGAGGAUGUUGUAUCUUUA. An siRNA targeting SNX3 was acquired from Dharmacon, target sequence: CGUGACUAUUAAUGAUUGA. A validated siRNA targeting PTPN23 was purchased from Thermo Fisher Scientific (s24775). The AllStars negative control siRNA was used as a non-specific control (Qiagen).

### Transfections

Plasmids: Cells were transfected 24 hours after plating, using Lipofectamine 2000, following the manufacturer’s protocol (Invitrogen). Lipofectamine 2000 was diluted in Opti-MEM (Gibco) and incubated for 5 minutes at RT prior to mixing with DNA diluted in Opti-MEM. The DNA:Lipofectamine mix was incubated for 20 minutes prior to addition to cells in antibiotic-free media. The cells were incubated with the transfection mix for 4 hours before the media was replaced.

### siRNAs

Cells were reverse-transfected (transfected while plating) in 6-well plates (3.5 -5 × 10^5^ cells per well) using Lipofectamine RNAimax transfection reagent (Invitrogen), following standard protocols. The siRNAs were diluted in Opti-MEM, and added to wells containing cells in antibiotic-free media at a 10 nM final concentration. Assays were performed 48 or 72 hours post-transfection with siRNAs, as indicated in figure legends.

### Flow Cytometry

To quantify cell-surface levels of CD4 and BST-2, HeLa cells transfected to express Vpu constructs or empty plasmid control, and the pCG-GFP transfection marker, were washed with 1X phosphate-buffered saline (PBS) and resuspended using Acutase dissociation media (Innovative Cell Technologies). The cells were collected and pelleted by centrifugation at 300 x g for 5 minutes, then resuspended in 100 µL flow cytometry buffer (2% FBS in PBS, and 0.1% sodium azide) and stained using either Alexa-647-conjugated mouse anti-BST-2 antibody, Alexa-647-IgG isotype control, APC-conjugated mouse anti-CD4, or APC-conjugated mouse IgG1 isotype control (BioLegend), and incubated for 30 minutes on ice. The cells were washed and pelleted three times before fixation in 2% paraformaldehyde (PFA) in 1X PBS for 15 minutes. Surface BST-2 or CD4 was quantified using a BD Accuri C6 flow cytometer and CFlow Sampler analysis software. Data are presented as mean fluorescence intensity of FL4 (far-red) signal in the GFP-positive (FL1) cell population.

### Immunofluorescence Microscopy

1.2 x 10^5^ HeLa P4.R5 cells were seeded on 12 mm coverslips in 24-well plates 24 hours prior to transfection. Cells were transfected with 200 ng DNA using Lipofectamine 2000, following the manufacturer’s protocol. The cells were fixed and stained the following day. The cells were washed in cold PBS and fixed in 4% PFA in PBS on ice for 5 minutes, then 15 minutes at room temperature (RT). The cells were washed twice with PBS and PFA was quenched with 50 mM ammonium chloride for 5 minutes. The cells were permeabilized with 0.2% Triton X-100 in 1X PBS for 7 minutes and blocked with 2% bovine serum albumin (BSA) for 30 minutes at RT prior to incubation with primary antibodies for 2 hours at RT. Vpu was detected with mouse anti-FLAG (Sigma-Aldrich); the *trans*-Golgi was detected with goat anti-TGN46 (ABD Serotec); Mito-V5-APEX2 was detected using mouse anti-V5 (Invitrogen); biotin was detected with Alexa-594-conjugated Streptavidin (Invitrogen). Endogenous protein cofactors were detected using rabbit anti-STAM antibody (ProteinTech), rabbit anti-PTPN23 (ProteinTech) goat anti-Vps35 (Novus Bio) and rabbit anti-SNX3 (Abcam).

The cells were washed and stained with donkey anti-mouse rhodamine-X (RhX) or donkey anti-sheep AlexaFluor-488 (Jackson ImmunoResearch) for 1 hour at RT. For detection of APEX2 biotinylation by immunofluorescence, the cells were incubated with 500 µM biotinyl-tyramide in pre-warmed medium for 30 minutes prior to addition of hydrogen peroxide (1 mM). The cells were incubated for 1 minute prior to quenching with APEX2 quencher solution (see below), and washing 3x with PBS before fixation and staining as above. Stretavidin conjugated to Alexa-Fluor 594 was used to detect biotinylated proteins. Following immunostaining, the cells were washed extensively in PBS, and briefly in water, before mounting in Mowiol (polyvinyl alcohol) mounting medium (prepared in-house).

Images were captured at 100x magnification (1344 × 1024 pixels) using an Olympus IX81 wide-field microscope fitted with a Hamamatsu CCD camera. For each field, a Z-series of images was collected, deconvolved using a nearest-neighbor algorithm (Slidebook software v6, Imaging Innovations, Inc) and presented as Z-stack projections. Image insets displaying colocalization are single Z-plane images. Image brightness was adjusted using Adobe Photoshop CS3.

### Transmission Electron Microscopy

6 x 10^5^ HeLa P4.R5 cells were seeded in 35 mm poly-lysine-coated MatTek dishes and transfected 24 hours later with 1 µg total pcDNA3.1-VpHu-FLAG-APEX2 or pcDNA3.1-VpHu-S52,56N-FLAG-APEX2. 16 hours later, the cells were fixed in 2% glutaraldehyde (Electron Microscopy Sciences) in 100 mM sodium cacodylate with 2 mM CaCl_2_, pH 7.4, for 60 minutes on ice. All subsequent steps were performed on ice until resin infiltration. The cells were rinsed with 100 mM sodium cacodylate with 2 mM CaCl_2_ five times for two minutes before addition of 20 mM glycine 100 mM sodium cacodylate with 2 mM CaCl_2_ to quench unreacted fixative. The cells were washed with 100 mM sodium cacodylate with 2 mM CaCl_2_ and diaminobenzidine (DAB) staining was initiated with the addition of freshly diluted 0.5 mg/mL DAB (Sigma; from a stock of the free base dissolved in 0.1 M HCl) and 0.03% H_2_O_2_ in 100 mM sodium cacodylate with 2 mM CaCl_2_. After 5 minutes, the reaction was stopped with the removal of the DAB solution, and the cells were again washed with 100 mM sodium cacodylate with 2 mM CaCl_2_. Post-fixation staining was performed with 2% (w/v) osmium tetroxide (Electron Microscopy Sciences) for 30 minutes in chilled buffer. Cells were rinsed 5× 2 minutes each in chilled distilled water and then placed in chilled 2% (w/v) uranyl acetate in ddH_2_O (Electron Microscopy Sciences) overnight. Cells were washed in distilled water, and dehydrated in graded ethanol series (20%, 50%, 75%, 90%, 95%, 100%, 100%, 100%), for 2 minutes each. The cells were brought to RT in 100% ethanol, and infiltrated with Durcapan ACM resin (Sigma-Aldrich) diluted in ethanol 1:1 for one hour. The cells were then infiltrated with 100% resin twice for one hour each before curing at 60°C for 48 hours. DAB positive cells were identified at low resolution by wide-field microscopy. 70-90 nm-thin sections were imaged using FEI-Tecnai G2 Spirit or JEOL 1200EX transmission electron microscopes operating at 80kV. Sample processing and imaging by electron microscopy was performed at the National Center for Microscopy and Imaging Research at UC San Diego.

### Western Blot

Cell monolayers were washed 3 times in ice-cold PBS and lysed in extraction buffer (0.5% Triton X-100, 150 mM NaCl, 25 mM KCl, 25 mM Tris, pH 7.4, 1 mM EDTA) supplemented with a protease inhibitor mixture (Roche Applied Science). Extracts were clarified by centrifugation (12,000 × g for 10 minutes at 4°C). The sample protein concentration was determined by Bradford assay (BD Biosciences) using standard protocols, and 10 µg denatured by boiling for 5 minutes in SDS sample buffer. Proteins in the extracts were resolved by SDS-PAGE using 12% or 4-15% gradient (BioRad) acrylamide gels, transferred to PVDF membranes, and probed by immunoblotting using mouse anti-Actin (Sigma-Aldrich), mouse anti-FLAG (Sigma-Aldrich), mouse anti-V5 (Invitrogen), STAM (ProteinTech), Vps35 (Novus Bio), SNX3 (Abcam), PTPN23 (ProteinTech), and horseradish peroxidase-conjugated goat anti-Mouse IgG (BioRad) or HRP-donkey anti-Rabbit IgG (BioRad) and Western Clarity detection reagent (BioRad). Apparent molecular mass was estimated using commercial protein standards (PageRulePlus, Thermo Scientific). Chemiluminescence was detected using a BioRad Chemi Doc imaging system and analyzed using BioRad Image Lab v5.1 software.

### Protein Biotinylation

HeLa P4.R5 cells were plated in 10 cm dishes at 3.2x10^6^ cells per dish. The following day, the cells were transfected with 12 µg plasmid DNA using Lipofectamine 2000 (Invitrogen/Thermo Fisher), following manufacturer’s guidelines. Biotinylation and protein harvest was performed 24 hours later, following established protocol (41). The cells were incubated with 500 µM biotinyl-tyramide in pre-warmed complete DMEM for 30 minutes at 37 °C. The biotinylation reaction was catalyzed by addition of 1 mM hydrogen peroxide to the culture media for 1 minute before quenching three times with APEX2 quenching solution (10 mM sodium ascorbate, 5 mM Trolox, and 10 mM sodium azide in 1 X PBS). The cells were scraped from the dishes with the final quencher wash into 15 mL Falcon tubes and pelleted by centrifugation at 300 x g for 5 minutes at 4°C. The cell pellets were lysed in 1ml RIPA buffer containing quenching components and protease inhibitor cocktail (41) for 5 minutes on ice. The lysates were briefly vortexed and nuclei pelleted by centrifugation at 15,000 x g for 10 minutes at 4°C. Protein content from the supernate was measured by Bradford protein assay (BioRad) and equal amounts incubated with streptavidin beads at 4°C overnight, while gently agitated. The following day, the beads were washed 2x with RIPA lysis buffer, and 1x with 2 M urea in 10 mM Tris-HCl (pH 8). The beads were washed again with RIPA lysis buffer, 2 x 1x PBS, and protein eluted from the beads in excess Biotin (200 mM NaCl, 50 mM Tris-HCl (pH 8), 2% SDS, 1 mM D-Biotin) at 70 °C for 30 minutes. An aliquot was stored for Western blot analysis, and remaining eluate processed for mass spectrometric analysis.

### Quantitative Mass Spectrometry

Quantitative MS analysis was performed as previously described (42) in the Collaborative Center for Multiplexed Proteomics in the Department of Pharmacology and the Skaggs School of Pharmacy and Pharmaceutical Sciences at UC San Diego. All quantitative mass spectrometry experiments were performed in biological duplicate. Protein disulfides were reduced with 5 mM DTT at 56°C for 30 minutes. Proteins were cooled on ice and alkylated with 15 mM iodoacetamide for 20 minutes at RT. Reduced/alkylated proteins were precipitated by addition of trichloroacetic acid (TCA) on ice for 10 min. Precipitated proteins were pelleted by centrifugation at 14,000 rpm for 5min, the supernatent was removed, protein resuspended in cold acetone, pelleted, and acetone wash repeated. Precipitated proteins were re-suspended in 1 M urea in 50 mM HEPES, pH 8.5 for proteolytic digestion. Proteins were first digested with LysC overnight at room temperature, then with trypsin for 6 hours at 37°C. Digestion was quenched by the addition of 10% trifluoroacetic acid (TFA), and peptides were desalted with C18 solid-phase extraction columns. Peptides were dried in a speed-vacuum concentrator, then re-suspended in 50% Acetonitrile/5% formic acid and quantified by BCA assay. A 50 μg aliquot was made for each sample for proteomic analysis. Protein samples were labeled with TMT10plex isobaric mass tag labeling reagents (Thermo Scientific), at a concentration of 20 μg/μL in dry acetonitrile. Lyophilized peptides were re-suspended in 50 μL 30% acetonitrile in 200 mM HEPES, pH 8.5 and 8 μL of the appropriate TMT reagent was added to each sample. The labeling reaction was conducted for 1 hour at RT, and then quenched by the addition of 9 μL of 5% hydroxylamine for 15 minutes at RT. Labeled samples were then acidified by adding 50 μL of 1% TFA. Differentially labeled samples were pooled into multiplex experiments and then desalted via solid-phase extraction.

Combined multiplexes were lyophilized and re-suspended in 5% formic acid/5% acetonitrile for identification and quantification by LC-MS2/MS3. All LC-MS2/MS3 experiments were performed on an Orbitrap Fusion mass spectrometer with an in-line Easy-nLC 1000 with chilled autosampler. Peptides were eluted with a linear gradient from 11 to 30% acetonitrile in 0.125% formic acid over 165 minutes at a flow rate of 300 nL/minute and heating the column to 60°C. Electrospray ionization was achieved by applying 2000V through a stainless-steel T-junction at the inlet of the column. Data were processed using the ProteomeDiscoverer 2.1.0.81 software package. Data were normalized as detailed previously (42). The data from the proteomic experiments have been uploaded to ProteomeXchange (PXD023713) through MassIVE (MSV000086733).

### Global Arrayed Protein Stability Analysis (GAPSA)

The GAPSA assay was performed as previously described (28). cDNA clones for approximately 160 genes were isolated from the Human ORFeome V8.1 Collection (Broad Institute). cDNA concentrations were normalized to 10 ng/μL prior to spotting in poly-D-lysine-coated 384 well plates. 20 ng cDNA encoding Vpu-FLAG, Vpu-S52,56N-FLAG or LacZ-FLAG control diluted in Opti-MEM was added to each well. Fugene6 (Promega) transfection reagent was added and incubated for 25 minutes at room temperature. 20 µL DMEM containing 6 x 10^4^ HEK293T cells was added to each well and subsequently incubated for 48 hours at 37°C, 5 % CO_2_. The cells were stained using an automated protocol. First, the plates were washed with PBS and cells fixed in 8% paraformaldehyde for 1 hour at room temperature. The cells were then washed with PBS, and permeabilized with 0.5% Triton X-100 in PBS for 10 minutes at room temperature. The cells were then washed with PBS and incubated with 6% BSA in PBS for 1 hour at room temperature to block non-specific antigen binding. The cells were then washed with PBS and incubated with mouse anti-V5 and rabbit anti-FLAG antibodies (1:250) in BSA-PBS for 1 hour at room temperature. The cells then were washed with PBS and incubated with goat anti-mouse Alexa 488, and goat anti-rabbit Alexa 568 (1:250 dilution) in BSA-PBS for 1 hour at room temperature. Nuclei were then stained with 2 µg/mL DAPI. The plates were imaged using the Opera QEHS High-Content Imaging System. The image output was analyzed using the Acapella High-Content Image Analysis Software (PerkinElmer) using a custom script. Data analysis was done as previously described (28).

### Data presentation and statistics

Figures were prepared using Adobe Creative Suite CS3; immunofluorescence images were adjusted using Adobe Photoshop and figures prepared using Adobe Illustrator software. Statistical analyses were performed using Graphpad Prism v5. Mass spectrometry data was analyzed using Microsoft excel 2016 and R Studio (R v.4.0.3). Significance for proteomics data was assessed by Student’s *t*-test; variance was assessed by an F-test to ensure the correct statistical assumptions were used. *p* values of *p* ≤ 0.05 were considered significant. Heatmaps were generated using Morpheus matrix visualization software (Broad institute, https://software.broadinstitute.org/morpheus); data were sorted by k-means clustering, the optimal number of gene clusters was determined by elbow estimation method. Gene Ontology (GO) analysis was performed using the Database for Annotation, Visualization and Integrated Discovery (DAVID) v6.8 (43, 44). Network analysis of protein subset k-means cluster 6 was performed using the STRING app for Cytoscape (v3.7.0) (45, 46).

## Acknowledgements

We thank Klaus Strebel for the original codon-optimized VpHu construct and pNL4-3ΔVpu; Alice Ting for APEX-related constructs; the NIH AIDS Reagent Program; and The Pendleton Charitable Trust. The work was supported by NIH grants R37AI081668 to JCG, R01 AI124843 and R01 AI127302 to SKC, and in part by UC San Diego Center for AIDS Research Developmental Awards to CAS, LP, and DG; an NIH-funded program (P30 AI036214). SL was supported by a research fellowship of the Deutsche Forschungsgemeinschaft (DFG) (Grant reference number 404687549). JMW was supported by NIH T32 GM007752 and T32 AR064194, JL was supported by NIH K12 GM06852.

## Supplementary Materials

Table S1: Mass Spectrometry Experiment 1: Comparison of Mito Matrix and Vpu WT and mutants

Table S2: Mass Spectrometry Experiment 2 and 3: Comparison of Vpu WT and mutants

Table S3: Gene Ontology network analysis of differentially-enriched Vpu-proximal protein subsets

---

**Figure S1.**
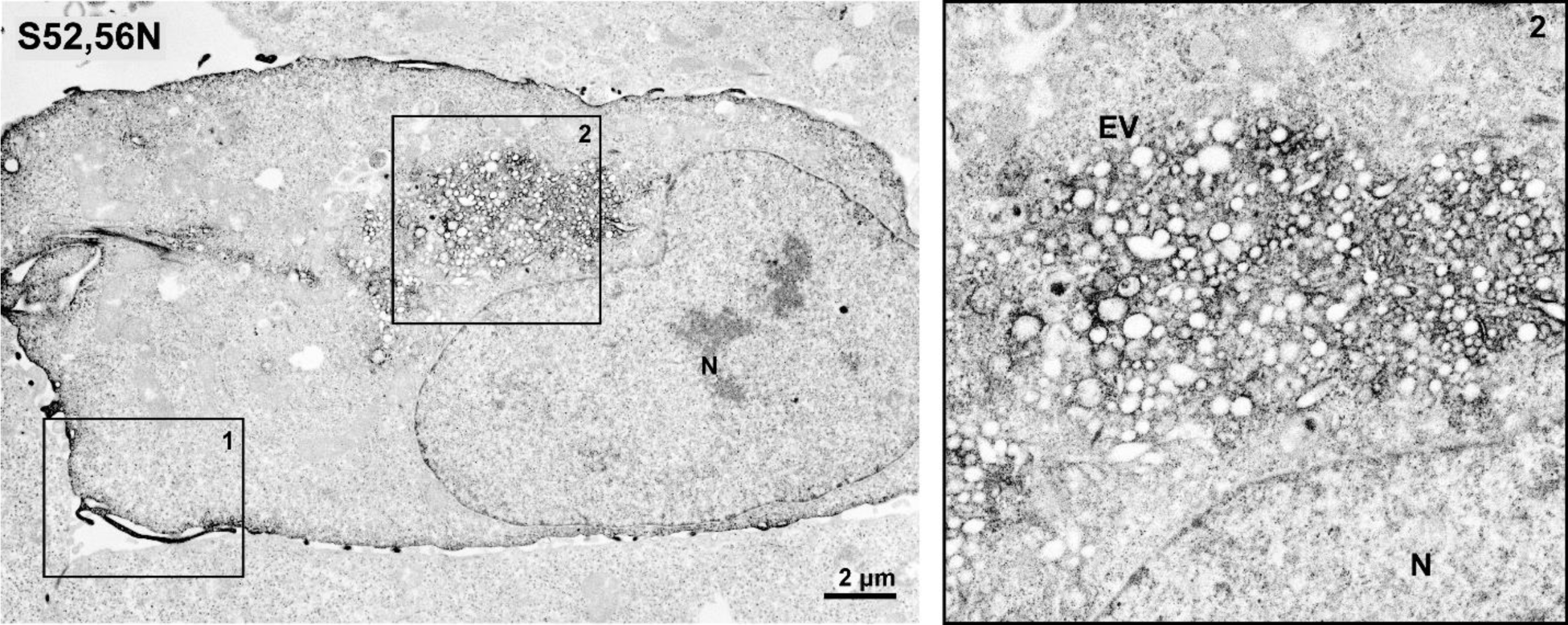
Juxtanuclear endosomal distortion in cells expressing Vpu-S52,56N-APEX2. HeLa P4.R5 cells were transfected to express Vpu-S52,56N-APEX2. 24 hours later the cells were fixed before APEX2-dependent polymerization of DAB and osmium staining. Cells were embedded in resin and 70 nm sections collected and analysed by TEM. The mutant Vpu-S52,56N was localized to the plasma membrane region (region 1 is shown at higher resolution in Fig. 3) but also induced formation of juxta-nuclear enlarged vesicles (EV, region 2), similar to the WT Vpu.

**Figure S2.**
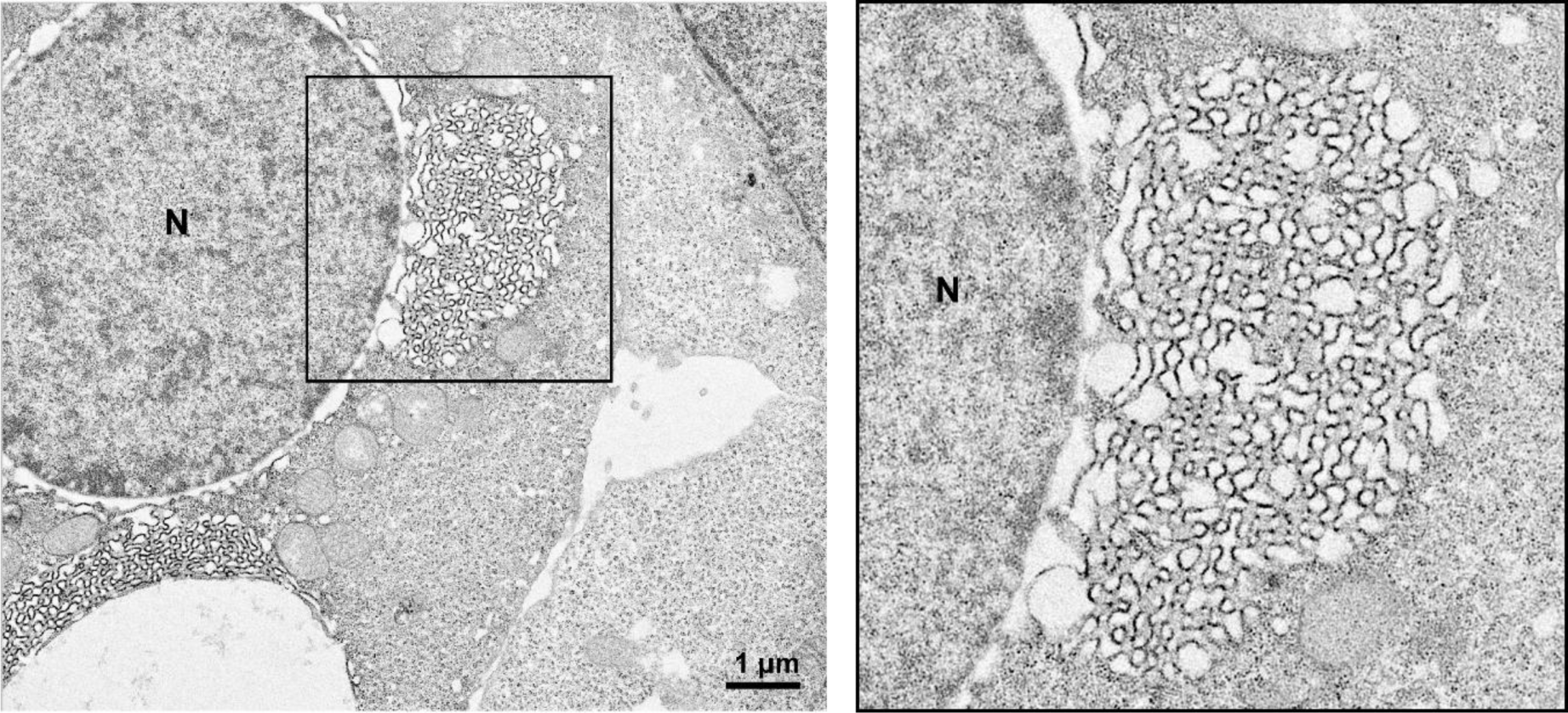
Exuberant ER membranes in cells expressing Vpu-A18H-APEX2. HeLa P4.R5 cells were transfected with a VpHu-A18H construct bearing a C-terminal APEX2 tag. 24 hours later the cells were fixed before APEX2-dependent polymerization of DAB and osmium staining. Cells were embedded in resin and 70 nm sections collected and analysed by TEM. The endoplasmic reticulum-trapped mutant, A18H, was restricted to the nuclear envelope (NE) and ER and induced membrane reorganisation: when expressed at high levels, the nuclear envelope was distorted by the accumulation of convoluted, smooth membranes.

**Figure S3.**
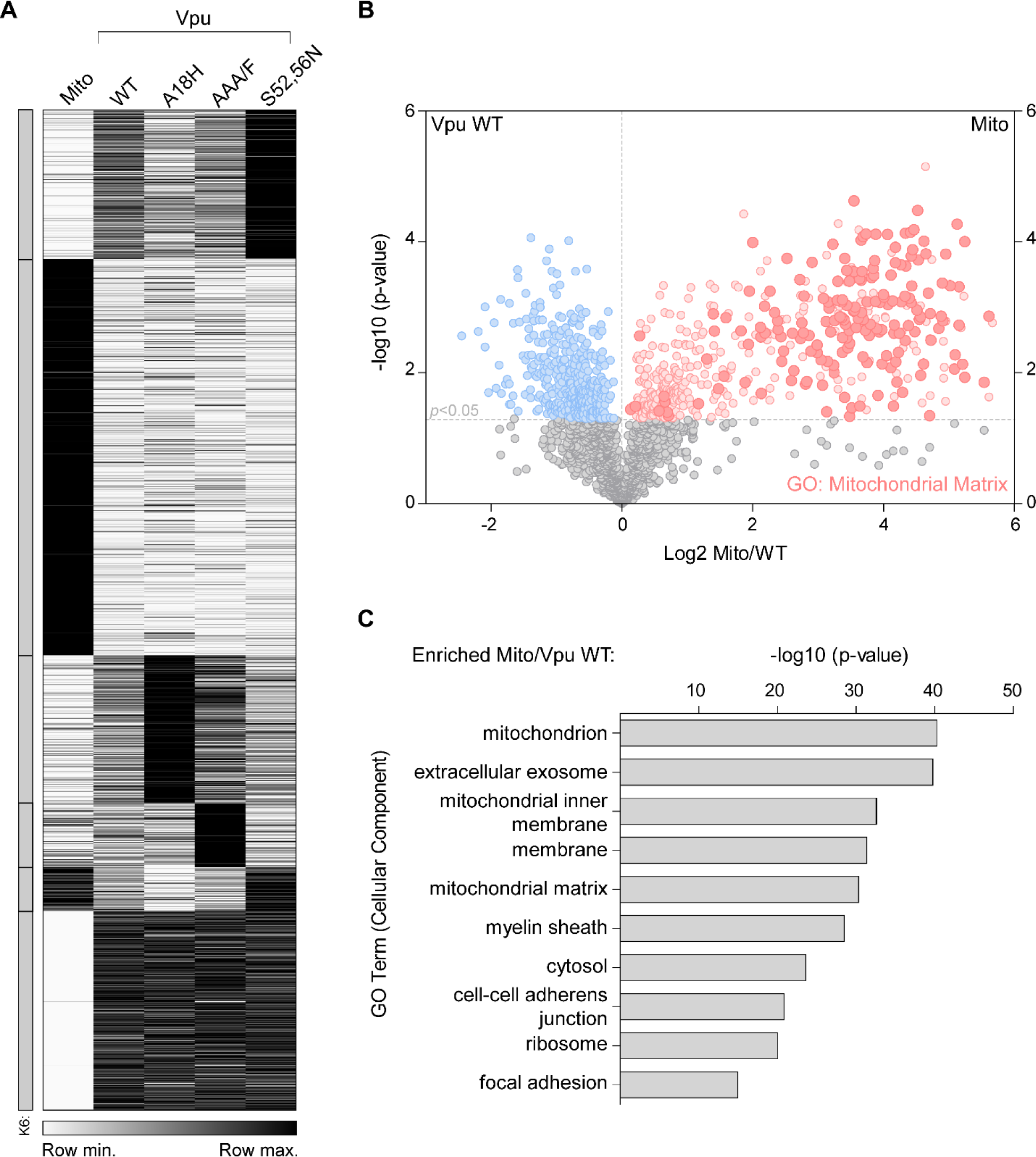
Pair-wise comparison of proximity-ome of Vpu-APEX2 compared to the Mito-APEX2. HeLa P4.R5 cells were transfected to express Mito (control) or Vpu constructs bearing C-terminal APEX2 tags, in duplicate. Following proximity biotinylation reactions, the biotinylated proteins were isolated and subject to quantitative mass spectrometry. (A) Heatmap showing relative protein abundance across Mito control and Vpu WT and mutant samples, sorted into 6 k-means clusters (cluster number derived from elbow method). (B) Volcano plot of proteins biotinylated by Vpu-APEX2 vs. Mito-APEX2 control. Mitochondrial proteins corresponding to GO term Mitochondrial Matrix are highlighted. The x-axis shows log2 fold change and y-axis -log10 *p*-value derived from Student’s *t*-test. (C) GO enrichment analysis of proteins significantly enriched by Mito-APEX compared to Vpu WT, the top ten GO (cell component) terms are shown.

**Figure S4.**
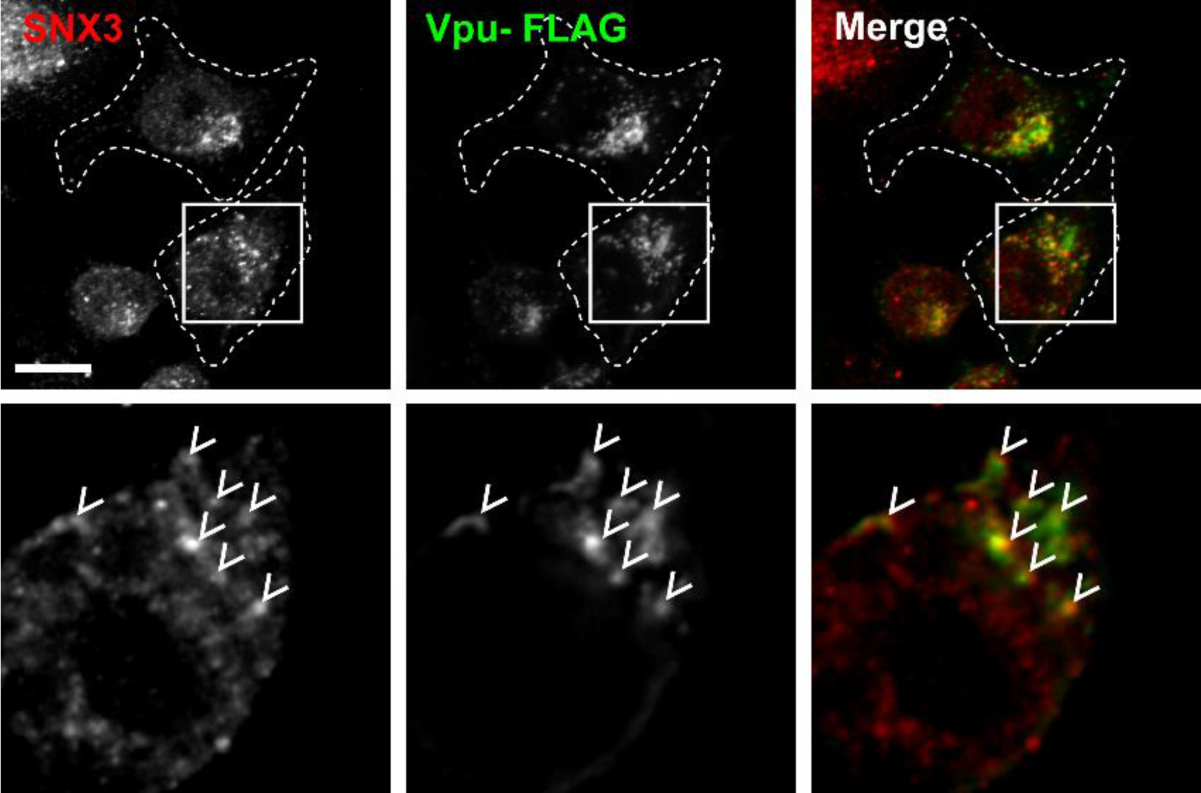
Immunofluorescence microscopy of Vpu-FLAG and candidate cofactor SNX3. HeLa P4.R5 cells were transfected to express Vpu-FLAG. Cells were fixed and stained for endogenous SNX3 protein 24 hours post-transfection. Images are z-stack projections of full cell volumes; insets show single z-sections, with arrows indicating colocalized foci. Scale bars are 10 µm. Some punctate colocalization of Vpu and SNX3 was observed in the perinuclear region, in agreement with immunofluorescent stain of Vpu and retromer component Vps35.

## References

1. Van Damme N, Goff D, Katsura C, Jorgenson RL, Mitchell R, Johnson MC, et al. The interferon-induced protein BST-2 restricts HIV-1 release and is downregulated from the cell surface by the viral Vpu protein. Cell host & microbe. 2008;3(4):245–52.

2. Neil SJ, Zang T, Bieniasz PD. Tetherin inhibits retrovirus release and is antagonized by HIV-1 Vpu. Nature. 2008;451(7177):425–30.

3. Lambele M, Koppensteiner H, Symeonides M, Roy NH, Chan J, Schindler M, et al. Vpu is the main determinant for tetraspanin downregulation in HIV-1-infected cells. J Virol. 2015;89(6):3247–55.

4. Willey RL, Maldarelli F, Martin MA, Strebel K. Human immunodeficiency virus type 1 Vpu protein induces rapid degradation of CD4. Journal of virology. 1992;66(12):7193–200.

5. Shah AH, Sowrirajan B, Davis ZB, Ward JP, Campbell EM, Planelles V, et al. Degranulation of natural killer cells following interaction with HIV-1-infected cells is hindered by downmodulation of NTB-A by Vpu. Cell host & microbe. 2010;8(5):397–409.

6. Ramirez PW, Famiglietti M, Sowrirajan B, DePaula-Silva AB, Rodesch C, Barker E, et al. Downmodulation of CCR7 by HIV-1 Vpu results in impaired migration and chemotactic signaling within CD4(+) T cells. Cell reports. 2014;7(6):2019–30.

7. Apps R, Del Prete GQ, Chatterjee P, Lara A, Brumme ZL, Brockman MA, et al. HIV-1 Vpu Mediates HLA-C Downregulation. Cell Host Microbe. 2016;19(5):686–95.

8. Margottin F, Bour SP, Durand H, Selig L, Benichou S, Richard V, et al. A novel human WD protein, h-beta TrCp, that interacts with HIV-1 Vpu connects CD4 to the ER degradation pathway through an F-box motif. Molecular cell. 1998;1(4):565–74.

9. Magadan JG, Perez-Victoria FJ, Sougrat R, Ye Y, Strebel K, Bonifacino JS. Multilayered mechanism of CD4 downregulation by HIV-1 Vpu involving distinct ER retention and ERAD targeting steps. PLoS pathogens. 2010;6(4):e1000869.

10. Mitchell RS, Katsura C, Skasko MA, Fitzpatrick K, Lau D, Ruiz A, et al. Vpu antagonizes BST-2-mediated restriction of HIV-1 release via beta-TrCP and endo-lysosomal trafficking. PLoS pathogens. 2009;5(5):e1000450.

11. Douglas JL, Viswanathan K, McCarroll MN, Gustin JK, Fruh K, Moses AV. Vpu directs the degradation of the human immunodeficiency virus restriction factor BST-2/Tetherin via a {beta}TrCP-dependent mechanism. Journal of virology. 2009;83(16):7931–47.

12. Schmidt S, Fritz JV, Bitzegeio J, Fackler OT, Keppler OT. HIV-1 Vpu blocks recycling and biosynthetic transport of the intrinsic immunity factor CD317/tetherin to overcome the virion release restriction. mBio. 2011;2(3):e00036–11.

13. Dube M, Roy BB, Guiot-Guillain P, Binette J, Mercier J, Chiasson A, et al. Antagonism of tetherin restriction of HIV-1 release by Vpu involves binding and sequestration of the restriction factor in a perinuclear compartment. PLoS pathogens. 2010;6(4):e1000856.

14. Lau D, Kwan W, Guatelli J. Role of the endocytic pathway in the counteraction of BST-2 by human lentiviral pathogens. Journal of virology. 2011;85(19):9834–46.

15. Kueck T, Neil SJ. A cytoplasmic tail determinant in HIV-1 Vpu mediates targeting of tetherin for endosomal degradation and counteracts interferon-induced restriction. PLoS pathogens. 2012;8(3):e1002609.

16. Stoneham CA, Singh R, Jia X, Xiong Y, Guatelli J. Endocytic Activity of HIV-1 Vpu: Phosphoserine-dependent Interactions with Clathrin Adaptors. Traffic. 2017.

17. Jia X, Weber E, Tokarev A, Lewinski M, Rizk M, Suarez M, et al. Structural basis of HIV-1 Vpu-mediated BST2 antagonism via hijacking of the clathrin adaptor protein complex 1. eLife. 2014;3:e02362.

18. Kueck T, Foster TL, Weinelt J, Sumner JC, Pickering S, Neil SJ. Serine Phosphorylation of HIV-1 Vpu and Its Binding to Tetherin Regulates Interaction with Clathrin Adaptors. PLoS pathogens. 2015;11(8):e1005141.

19. Janvier K, Pelchen-Matthews A, Renaud JB, Caillet M, Marsh M, Berlioz-Torrent C. The ESCRT-0 component HRS is required for HIV-1 Vpu-mediated BST-2/tetherin down-regulation. PLoS Pathog. 2011;7(2):e1001265.

20. Van Damme N, Guatelli J. HIV-1 Vpu inhibits accumulation of the envelope glycoprotein within clathrin-coated, Gag-containing endosomes. Cell Microbiol. 2008;10(5):1040–57.

21. Varthakavi V, Smith RM, Martin KL, Derdowski A, Lapierre LA, Goldenring JR, et al. The pericentriolar recycling endosome plays a key role in Vpu-mediated enhancement of HIV-1 particle release. Traffic. 2006;7(3):298–307.

22. Dube M, Roy BB, Guiot-Guillain P, Mercier J, Binette J, Leung G, et al. Suppression of Tetherin-restricting activity upon human immunodeficiency virus type 1 particle release correlates with localization of Vpu in the trans-Golgi network. Journal of virology. 2009;83(9):4574–90.

23. Lam SS, Martell JD, Kamer KJ, Deerinck TJ, Ellisman MH, Mootha VK, et al. Directed evolution of APEX2 for electron microscopy and proximity labeling. Nat Methods. 2015;12(1):51–4.

24. Martell JD, Deerinck TJ, Sancak Y, Poulos TL, Mootha VK, Sosinsky GE, et al. Engineered ascorbate peroxidase as a genetically encoded reporter for electron microscopy. Nature biotechnology. 2012;30(11):1143–8.

25. Hauser H, Lopez LA, Yang SJ, Oldenburg JE, Exline CM, Guatelli JC, et al. HIV-1 Vpu and HIV-2 Env counteract BST-2/tetherin by sequestration in a perinuclear compartment. Retrovirology. 2010;7:51.

26. Skasko M, Tokarev A, Chen CC, Fischer WB, Pillai SK, Guatelli J. BST-2 is rapidly down-regulated from the cell surface by the HIV-1 protein Vpu: evidence for a post-ER mechanism of Vpu-action. Virology. 2011;411(1):65–77.

27. Skasko M, Wang Y, Tian Y, Tokarev A, Munguia J, Ruiz A, et al. HIV-1 Vpu protein antagonizes innate restriction factor BST-2 via lipid-embedded helix-helix interactions. The Journal of biological chemistry. 2012;287(1):58–67.

28. Jain P, Boso G, Langer S, Soonthornvacharin S, De Jesus PD, Nguyen Q, et al. Large-Scale Arrayed Analysis of Protein Degradation Reveals Cellular Targets for HIV-1 Vpu. Cell reports. 2018;22(9):2493–503.

29. Doyotte A, Mironov A, McKenzie E, Woodman P. The Bro1-related protein HD-PTP/PTPN23 is required for endosomal cargo sorting and multivesicular body morphogenesis. Proc Natl Acad Sci U S A. 2008;105(17):6308–13.

30. Matheson NJ, Sumner J, Wals K, Rapiteanu R, Weekes MP, Vigan R, et al. Cell Surface Proteomic Map of HIV Infection Reveals Antagonism of Amino Acid Metabolism by Vpu and Nef. Cell host & microbe. 2015;18(4):409–23.

31. Jager S, Cimermancic P, Gulbahce N, Johnson JR, McGovern KE, Clarke SC, et al. Global landscape of HIV-human protein complexes. Nature. 2011;481(7381):365–70.

32. Ali N, Zhang L, Taylor S, Mironov A, Urbe S, Woodman P. Recruitment of UBPY and ESCRT exchange drive HD-PTP-dependent sorting of EGFR to the MVB. Curr Biol. 2013;23(6):453–61.

33. Raiborg C, Bache KG, Gillooly DJ, Madshus IH, Stang E, Stenmark H. Hrs sorts ubiquitinated proteins into clathrin-coated microdomains of early endosomes. Nat Cell Biol. 2002;4(5):394–8.

34. Barlowe C, Schekman R. SEC12 encodes a guanine-nucleotide-exchange factor essential for transport vesicle budding from the ER. Nature. 1993;365(6444):347–9.

35. Dube M, Paquay C, Roy BB, Bego MG, Mercier J, Cohen EA. HIV-1 Vpu antagonizes BST-2 by interfering mainly with the trafficking of newly synthesized BST-2 to the cell surface. Traffic. 2011;12(12):1714–29.

36. Charneau P, Mirambeau G, Roux P, Paulous S, Buc H, Clavel F. HIV-1 reverse transcription. A termination step at the center of the genome. Journal of molecular biology. 1994;241(5):651–62.

37. Greenberg ME, Bronson S, Lock M, Neumann M, Pavlakis GN, Skowronski J. Co-localization of HIV-1 Nef with the AP-2 adaptor protein complex correlates with Nef-induced CD4 down-regulation. The EMBO journal. 1997;16(23):6964–76.

38. Yang X, Boehm JS, Yang X, Salehi-Ashtiani K, Hao T, Shen Y, et al. A public genome-scale lentiviral expression library of human ORFs. Nature methods. 2011;8(8):659–61.

39. Adachi A, Gendelman HE, Koenig S, Folks T, Willey R, Rabson A, et al. Production of acquired immunodeficiency syndrome-associated retrovirus in human and nonhuman cells transfected with an infectious molecular clone. Journal of virology. 1986;59(2):284–91.

40. Strebel K, Klimkait T, Martin MA. A novel gene of HIV-1, vpu, and its 16-kilodalton product. Science. 1988;241(4870):1221–3.

41. Hung V, Udeshi ND, Lam SS, Loh KH, Cox KJ, Pedram K, et al. Spatially resolved proteomic mapping in living cells with the engineered peroxidase APEX2. Nature protocols. 2016;11(3):456–75.

42. Lapek JD, Jr., Lewinski MK, Wozniak JM, Guatelli J, Gonzalez DJ. Quantitative Temporal Viromics of an Inducible HIV-1 Model Yields Insight to Global Host Targets and Phospho-Dynamics Associated with Protein Vpr. Mol Cell Proteomics. 2017;16(8):1447–61.

43. Huang da W, Sherman BT, Lempicki RA. Systematic and integrative analysis of large gene lists using DAVID bioinformatics resources. Nature protocols. 2009;4(1):44–57.

44. Huang da W, Sherman BT, Lempicki RA. Bioinformatics enrichment tools: paths toward the comprehensive functional analysis of large gene lists. Nucleic acids research. 2009;37(1):1–13.

45. Shannon P, Markiel A, Ozier O, Baliga NS, Wang JT, Ramage D, et al. Cytoscape: a software environment for integrated models of biomolecular interaction networks. Genome Res. 2003;13(11):2498–504.

46. Szklarczyk D, Gable AL, Lyon D, Junge A, Wyder S, Huerta-Cepas J, et al. STRING v11: protein-protein association networks with increased coverage, supporting functional discovery in genome-wide experimental datasets. Nucleic acids research. 2019;47(D1):D607–D13.

